# Membrane phase separation drives organization at B cell receptor clusters

**DOI:** 10.1101/2021.05.12.443834

**Authors:** Sarah A. Shelby, Ivan Castello-Serrano, Kathleen C. Wisser, Ilya Levental, Sarah L. Veatch

## Abstract

Heterogeneity in intact cell plasma membranes has been explained by analogy to coexisting liquid-ordered and liquid-disordered phases, although models based on this idea fall short of describing the rich structure within cell membranes. Here, a new framework of lipid-driven plasma membrane heterogeneity is presented, drawing on quantitative measurements of protein partitioning and dynamics within B cell receptor clusters in live B lymphocyte plasma membranes, compared to coexisting phases in isolated plasma membranes. We propose that membrane domains in cells integrate the thermodynamic state of the membrane and the magnitude of the applied stimulus to give rise to a tunable response. This framework is supported through functional observations of B cell receptor phosphorylation in perturbed systems.

## INTRODUCTION

The impact of phase separation within cells has become appreciated in recent years, largely through visualizations of micron-scale, high-contrast protein- and/or nucleic acid-rich liquid droplets in the cytoplasm and nucleus (Lyon et al., 2021; Shin and Brangwynne, 2017). Analogous phase separation has long been proposed to be an organizing principle in cell membranes, especially within the plasma membrane, where domains are often referred to as “lipid rafts” (Brown and Rose, 1992; Simons and Ikonen, 1997). Membrane domains were first proposed to explain functional observations in cells, and decades of work links lipid rafts to all facets of plasma membrane function, including trafficking, polarization, sensing, and transport (Dart, 2010; Delos Santos et al., 2015; Hanzal-Bayer and Hancock, 2007; Igarashi et al., 2020; Lingwood and Simons, 2010; Strzyz, 2019). The evidence for a role for lipid rafts in immune receptor signaling is particularly strong and longstanding, with past work evoking domains in the initial activation of receptors and the modulation of signaling responses (Holowka and Baird, 2016; Pierce and Liu, 2010; Varshney et al., 2016). This past work proposes that clustered receptors translocate into rafts, sequestering them from downregulating phosphatases while recruiting activating kinases.

While phase separation has emerged as a likely mechanism underlying functional heterogeneity in cell membranes (Eggeling et al., 2009; Koyama-Honda et al., 2020; Sanchez et al., 2012; Sengupta et al., 2007), it is clear that macroscopic phase separation does not occur, and models used to rationalize rafts as phase separated domains either lack utility or are easily disproved, leading to valid skepticism (Kenworthy, 2008; Kraft, 2013; Munro, 2003). In contrast, plasma membranes isolated from cells clearly separate into coexisting liquid-ordered (Lo) and liquid-disordered (Ld) phases (Baumgart et al., 2007), there is clear evidence that membrane composition is carefully tuned and actively maintained by the cell (Ernst et al., 2016; Sviridov and Miller, 2020), and there are examples where this biological tuning impacts the membrane phase transition (Burns et al., 2017; Cammarota et al., 2020; Levental et al., 2020b). A goal of the current work is to address the disconnect between the apparent importance of the phase transition with abundant observations indicating that the membrane is in a single phase under physiological conditions, drawing from an increasingly sophisticated picture of the thermodynamics of phase separation in membranes (Levental et al., 2020a; Shaw et al., 2021). Specifically, several proposed physical models explain how mechanisms rooted in phase separation lead to structure in membranes consistent with rafts as small and dynamic membrane domains or larger domains scaffolded by membrane proteins (Schmid, 2017). Some models show that membranes without macroscopic phase separated domains can still retain some properties of phase separated systems, raising the possibility of a connection between phase separation, “lipid rafts”, and the impact of membrane composition on biological function.

Here, we test if concepts of phase separation translate to intact and living cell membranes using single molecule fluorescence localization microscopy in B cells. Our methods are sensitive and quantitative, and exploit the nanoscale signaling platforms that assemble following B cell receptor (BCR) activation to template long lived membrane domains (Gold and Reth, 2019; Gupta and DeFranco, 2007; Pierce and Liu, 2010; Sohn et al., 2006; Stone et al., 2017). The thermodynamics of phase separation dictates the composition of coexisting phases, therefore we directly measure the concentration of probe molecules in the vicinity of BCR clusters at high spatial resolution. We interrogate numerous membrane probes with varying structures and topologies to isolate a generic contribution from phase separation. With this extensive dataset, we demonstrate that anchor phase preference is a quantitative determinant of its local concentration and dynamics within BCR signaling platforms, and that manipulation of phase-driven partitioning impacts BCR function. This convergence between live cell and equilibrated vesicle measurements directly and conclusively establishes phase separation as a functional organizing principle in cell plasma membranes.

## RESULTS

### BCR clusters mark stabilized domains in the plasma membranes of B cells

Figure 1 shows CH27 B cells expressing the fluorophore mEos3.2 anchored to the plasma membrane via similar yet biophysically distinct lipidated peptide motifs. One motif contains myristoyl and palmitoyl groups (PM) and the second contains a myristoyl modification and a polybasic sequence (M). Both are derived from Src family kinases that anchor to the inner leaflet of the plasma membrane. These probes are imaged in live cells alongside BCR labeled with an antibody fragment (Fab) against the μ subunit of IgM BCR that is both biotinylated and fluorescently tagged with an organic fluorophore (silicon rhodamine, SiR). SiR and mEos3.2 probes are imaged in cells under conditions where probes are isolated and can be localized, and reconstructed images show the positions of probes acquired over time (Fig 1A). BCR crosslinking organizes receptors into tight clusters (Pierce and Liu, 2010), producing reconstructed images in which puncta decorate the cell surface while both PM and M anchors are uniformly distributed. Both single molecules and BCR puncta are dynamic. Supplementary Movies 1-6 show single molecule motions and the evolution of BCR clusters over time.

**Figure 1:**
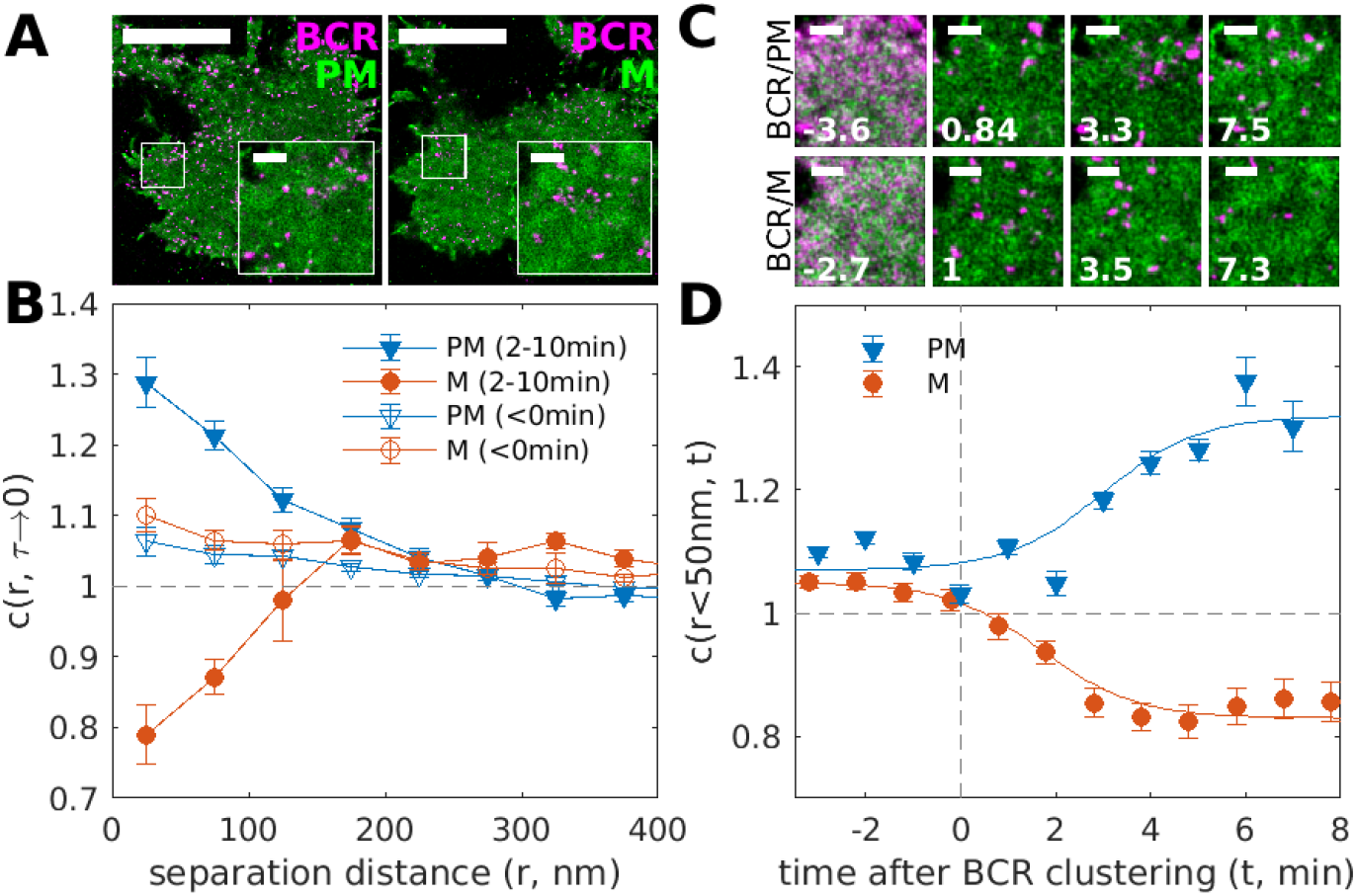
BCR clustering produces membrane domains that differently sort two fluorescent proteins anchored at the inner leaflet. (A) Reconstructed live-cell super-resolution images of CH27 B cells imaged between 2 and 10 min after BCR clustering. B cells expressing mEos3.2 conjugated to either a palmitoylated and myristoylated anchor (PM) or a myristoylated anchor (M) (green) are labeled with biotinylated and SiR labeled Fab anti-IgMμ (magenta) that is crosslinked with soluble streptavidin. (B) Quantification of BCR-anchor co-distributions both before (<0min) and after (2-10min) streptavidin addition via the spatial-temporal cross-correlation function c(r, τ) extrapolated to τ=0. Values of *c*(*r, τ* → 0) = 1 indicate a random co-distribution, *c*(*r,τ* → 0) > 1 indicate co-clustering, and *c*(*r,τ* → 0) < 1 indicate exclusion. (C) Images of the region of interest from A reconstructed from 2min of acquired data centered at the time indicated. Time-lapse movies for these cells showing single molecule motions or reconstructed images are included as Supplementary Movies 1-6. (D) The value of the *c*(*r,τ* → 0) in the first spatial bin (r<50nm) as a function of stimulation time averaged over 5 cells each expressing either PM or M. Cross-correlation amplitudes are evaluated using 2min of acquired localizations centered at the time indicated, as in C. Lines are drawn to guide the eye and are not a fit to any theory. Scale-bars are 10μm in A and are 1μm in A insets and C.

We have previously developed analytical methods for quantifying co-localization in super-resolved dynamic systems using spatio-temporal cross-correlation (Stone and Veatch, 2015). Cross-correlation functions are histograms of the pairwise distances between localized single molecules observed in different color channels, normalized by the frequency of expected pairwise-distances in a randomized co-distribution within the same region of interest and time-separation. To obtain statistical significance, cross-correlation curves are generated over an extended time-window (Δt=minutes), although we restrict our analysis to pairs observed at short separation-times (*τ* → 0) to extract accurate colocalization information of mobile components. Cross-correlation histograms are generated with spatial bins of 50nm, such that the first point in all curves is drawn at the bin center of 25nm and reflects the frequency of pairs with separation distances less than 50nm (r<50nm). See Materials and Methods and Supplementary Figure 1 for methodological details.

The cross-correlation functions of Fig 1B *c*(*r*, *τ* → 0) quantify the differential partitioning of PM and M anchors with respect to clustered BCR (closed symbols) and are tabulated from the same localizations used to generate the reconstructed images of Fig 1A. The curve for PM takes on values larger than 1 at short separation distances, then decays to 1 as the separation distance increases. The value of *c*(*r* < 50*nm*, τ *→* 0) = 1.3 indicates that the local density of PM within 50nm of the average BCR is 30% greater than the average density in the membrane overall. The inverse behavior is observed for the M anchor, with *c*(*r* < 50*nm*, *τ* → 0) = 0.8 indicating that M is 20% depleted within 50nm of the average BCR compared to the membrane overall. Cross-correlation functions tabulated from data acquired prior to crosslinking BCR do not show the same differential partitioning (open symbols). Both PM and M anchors are weakly correlated with uncross-linked BCR.

Figures 1C-D show the results of a similar analysis using a shorter time window Δt to better visualize how anchor organization evolves in time. The images in Fig 1C show insets from the same cells as in Fig 1A, but here images are reconstructed from localizations acquired within Δt=2min windows. Fig 1D shows how the anchor co-localization with BCR evolves with time, focusing only on cross-correlations at short separations from BCR (r<50nm) and averaging over several cells to improve statistics. Prior to crosslinking, BCR is uniformly distributed and is weakly correlated with both of the two anchors. After streptavidin crosslinking, receptors assemble into tight puncta and differential cross-correlations with mEos3.2-anchored peptides emerge over a timescale of several minutes. These results indicate that BCR clusters differentially sort probes distinguished only by the structure of their membrane anchors.

### Differential partitioning at BCR clusters extends across a panel of plasma membrane anchors

We extended these analyses to a total of 16 plasma membrane probes (presented schematically in Figure 2A; sequences are included in Supplementary Table 1) representing different modes of membrane anchorage. Many of these probes are truncated anchors from endogenous proteins, while others represent engineered constructs. Representative reconstructed images of these anchors alongside BCR imaged from 2-10 min after BCR crosslinking are shown in Fig 2A, along with curves showing the change in c(r) induced by BCR clustering and cellular activation (Δc(r)). Fig 2B summarizes the magnitude of crosslinking-induced changes in c(r) across all anchors at the smallest separation distances (r<50nm) (see Supplementary Figures S2-S3 for raw cross-correlation curves, summary histograms, and distributions of individual cell values).

**Figure 2:**
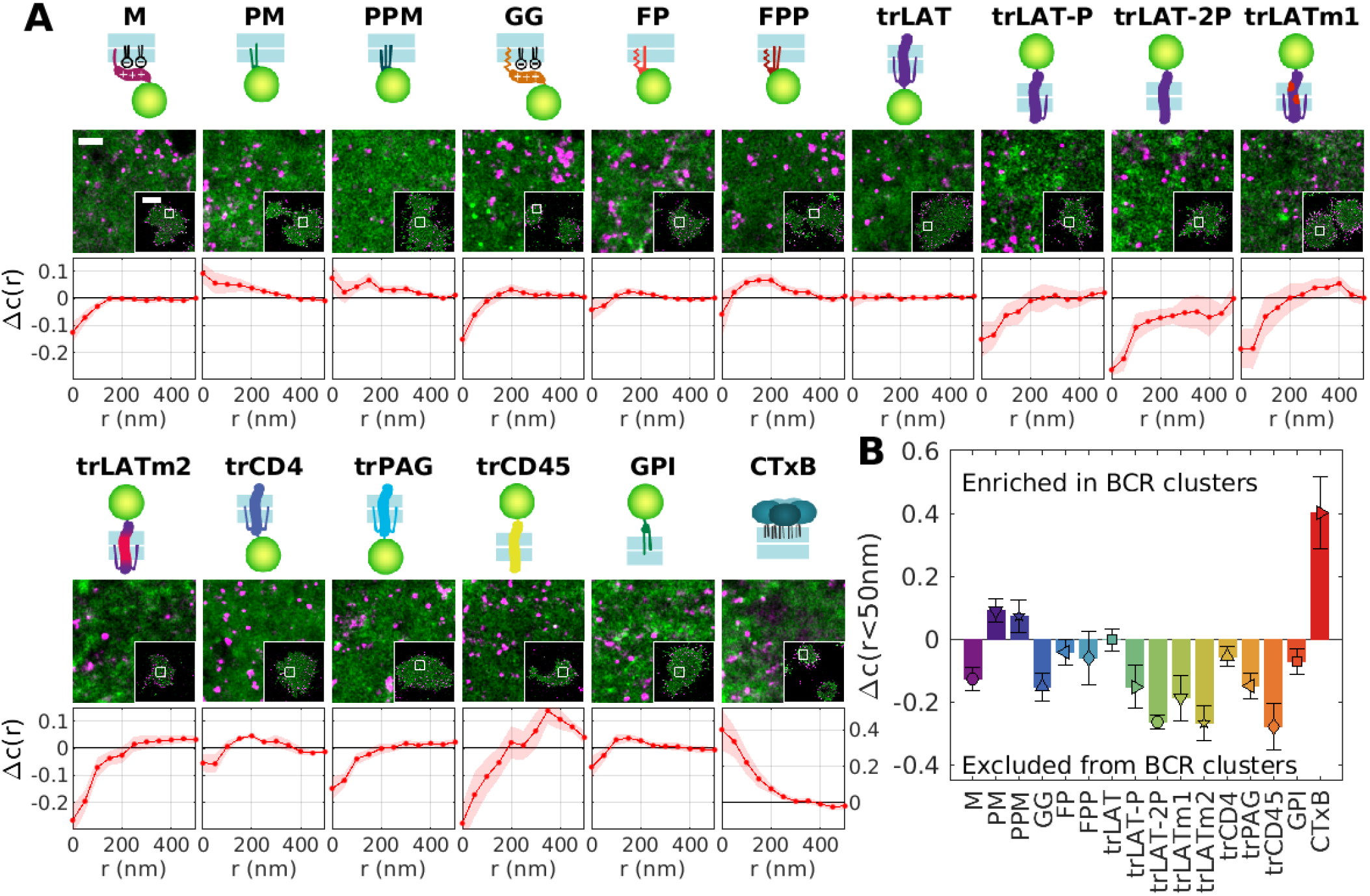
Anchored probes differentially partition with respect to clustered B cell receptors in live cells. (A) Schematic representation of anchors used in this study alongside representative reconstructed images showing anchor fluorescence (green) and BCR (magenta). The trace shows the change in c(r) between anchors and BCR for times between 2 and 10 min after BCR clustering and the curve obtained for times before BCR clustering, which we call Δc(r). Error bounds indicated by the shaded region represent the SEM over cells. (B) Summary of Δc(r) for separation distances less than 50nm. Δc(r<50nm) =0 indicates the anchor does not change its co-distribution with respect to BCR when BCR is clustered, Δc(r<50nm) >0 indicates anchors become more enriched near BCR after clustering, and Δc(r<50nm) <0 indicates anchors become more depleted near BCR after clustering.

The first class of probes are peripheral and anchored to the plasma membrane inner leaflet via lipidation, sometimes in combination with a polybasic sequence. In addition to M and PM anchors, this class includes the anchor PPM with a myristoylation and two palmitoylation sites derived from the Src family kinase Fyn, along with 3 anchors from the Ras family GTPases consisting of a prenyl (geranylgeranyl) group and a polybasic sequence (GG), a farnesyl group and a single palmitoylation (FP), or a farnesyl group and two palmitoylations (FPP). Across this series, anchors lacking palmitoyl groups (M and GG) become robustly excluded from BCR clusters, while palmitoylated anchors become enriched (PM or PPM) or remain neutrally distributed (FP and FPP) within 50 nm of clustered BCR. The second class of membrane anchors are single-pass transmembrane peptides. These include several variants of the transmembrane helix of the Linker for activation of T cells (“transmembrane LAT”; trLAT), where either transmembrane residues or palmitoylation sites were modified. Also included are minimal transmembrane anchor sequences of CD4, PAG/CBP, and CD45 (trCD4, trPAG, and trCD45). Most transmembrane anchors become excluded from BCR clusters to differing degrees, with the exception of trLAT and trCD4 which are both palmitoylated. We also examined two probes that localize exclusively to the outer leaflet of the plasma membrane, including a minimal GPI-linked protein anchor and exogenous cholera toxin subunit B (CTxB). CTxB is highly enriched at BCR clusters, while GPI is somewhat depleted at short separation distances.

Several anchors produce Δc(r) curves with local maxima away from r<50nm, suggesting that these anchors are maximally enriched at and beyond the boundaries of BCR clusters after B cell activation, which are estimated to have radii of 33±4nm (Supplementary Figure S4). This is observed for both FP and FPP inner leaflet anchors, trCD4 of the transmembrane anchors, and GPI of the extracellular anchors (Supplementary Figure S5). We speculate that this could reflect the asymmetric nature of some anchors that contain individual structural elements that may exhibit preferences for different local environments (e.g. prenylation and palmitoylation for FP and FPP). Another possibility is that the center of BCR clusters may represent a crowded environment and some anchors may be excluded due to steric occlusion (possibly trCD4 and/or GPI).

### Differential partitioning of anchors to the Lo phase in GPMVs

The anchors described in Fig 2 also exhibit diverse partitioning behavior in isolated giant plasma membrane vesicles (GPMVs) with coexisting liquid-ordered (Lo) and liquid-disordered (Ld) phases (Figure 3). To quantify this effect, GPMVs were prepared from cells expressing anchors under conditions shown previously to preserve protein palmitoylation (Levental et al., 2010). GPMVs were imaged using epi-fluorescence microscopy under conditions where vesicles robustly separate into coexisting Lo and Ld phases, visualized by a fluorescent lipid analog with strong preference for the Ld phase. Representative images of isolated GPMVs with M and PM anchors are shown in Fig 3A, and representative images for all anchors are included in Supplementary Figure S6. Phase partitioning is quantified by measuring the background-corrected fluorescence intensity of anchors in the two phases, which is proportional to their concentration. Anchor enrichment in the Lo phase is calculated from line-scans through intensity images of the vesicle mid-plane (as in Fig 3A) according to: ([Lo]-[Ld])/([Lo]+[Ld]). This quantity is bounded by −1 for an anchor that is exclusively present in the Ld phase and 1 for probes exclusively present in the Lo phase.

**Figure 3:**
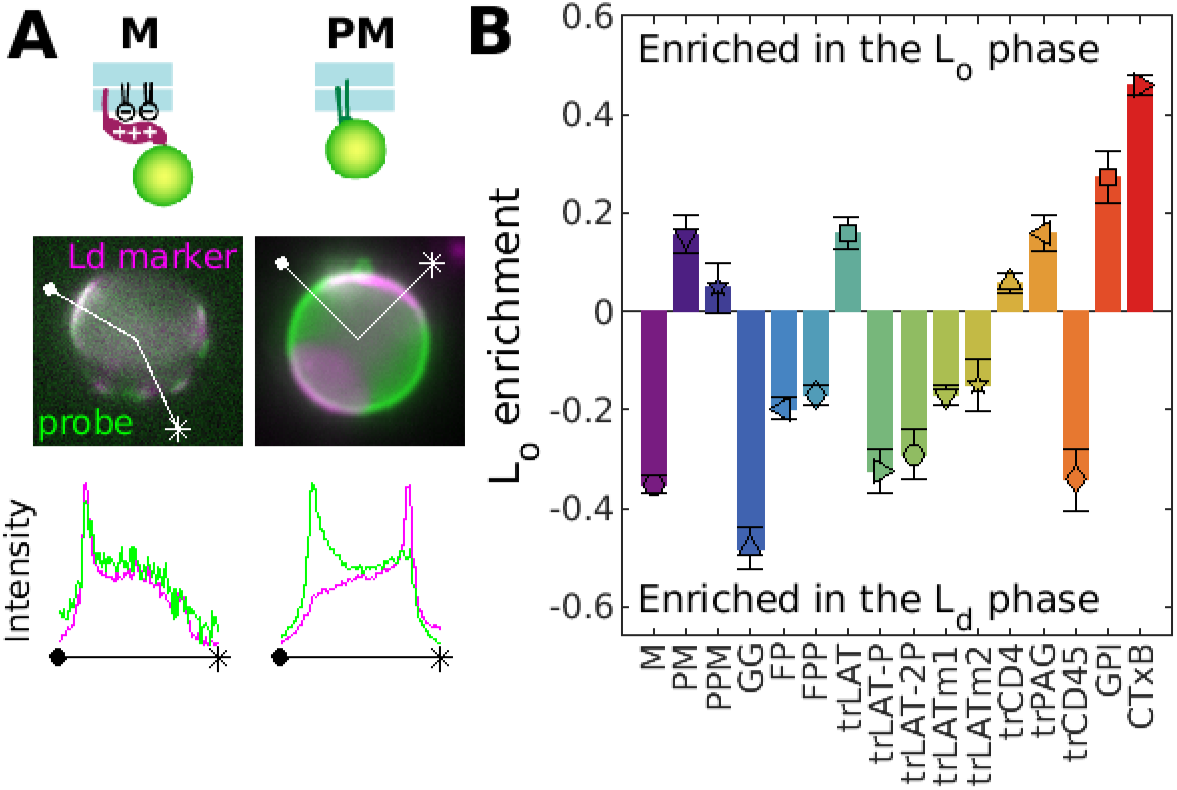
Anchored probes differentially partition with respect to Lo and Ld phases in isolated GPMVs. (A) Schematic representation of anchors from Fig 1 with fluorescence images of phase separated GPMVs showing anchor fluorescence (green) alongside a fluorescent marker for the Ld phase (DiO or DiI, magenta). Line scans show the fluorescence intensity in both channels along the trajectory shown. Similar images for all anchored probes are shown in Supplementary Figure S6. (B) Quantification of the Lo phase enrichment of anchors. Error bars show SEM over multiple vesicles and experiments. At least 10 vesicles across at least 3 separate experiments are used to produce the values shown in B.

The trends observed across anchors are in good qualitative agreement with expectations from past findings (Diaz-Rohrer et al., 2014; Levental et al., 2010; Lorent et al., 2017). In particular, palmitoylation is required but not sufficient for enrichment in the Lo phase for both transmembrane and inner leaflet peripheral anchors. Anchors that are both palmitoylated and myristoylated are more strongly enriched in Lo domains than anchors that are palmitoylated and prenylated. For transmembrane anchors, both palmitoylation and the sequence of the transmembrane helix impact the magnitude of Lo phase enrichment. GPI and CTxB anchors that localize exclusively to the outer leaflet both enrich strongly in the Lo phase.

### Probe concentration in BCR clusters is quantitatively predicted by GPMV phase partitioning

Figure 4A directly compares anchor partitioning near BCR clusters in live B cells (Δc(r<50nm)) with Lo enrichment of anchors in isolated GPMVs (Fig 3B). There is a statistically significant correlation between anchor partitioning in these two contexts when all anchors are considered together (p=0.002; Fig 4A). Here, the scatter of the points around the linear trend is larger than expected given the error bounds of individual measurements, indicating that multiple factors including Lo phase enrichment contribute to anchor co-localization with BCR clusters. Similar plots showing probe enrichment in cells with unclustered or clustered BCR (c(r<50nm)) are included in Supplementary Figure S7.

**Figure 4:**
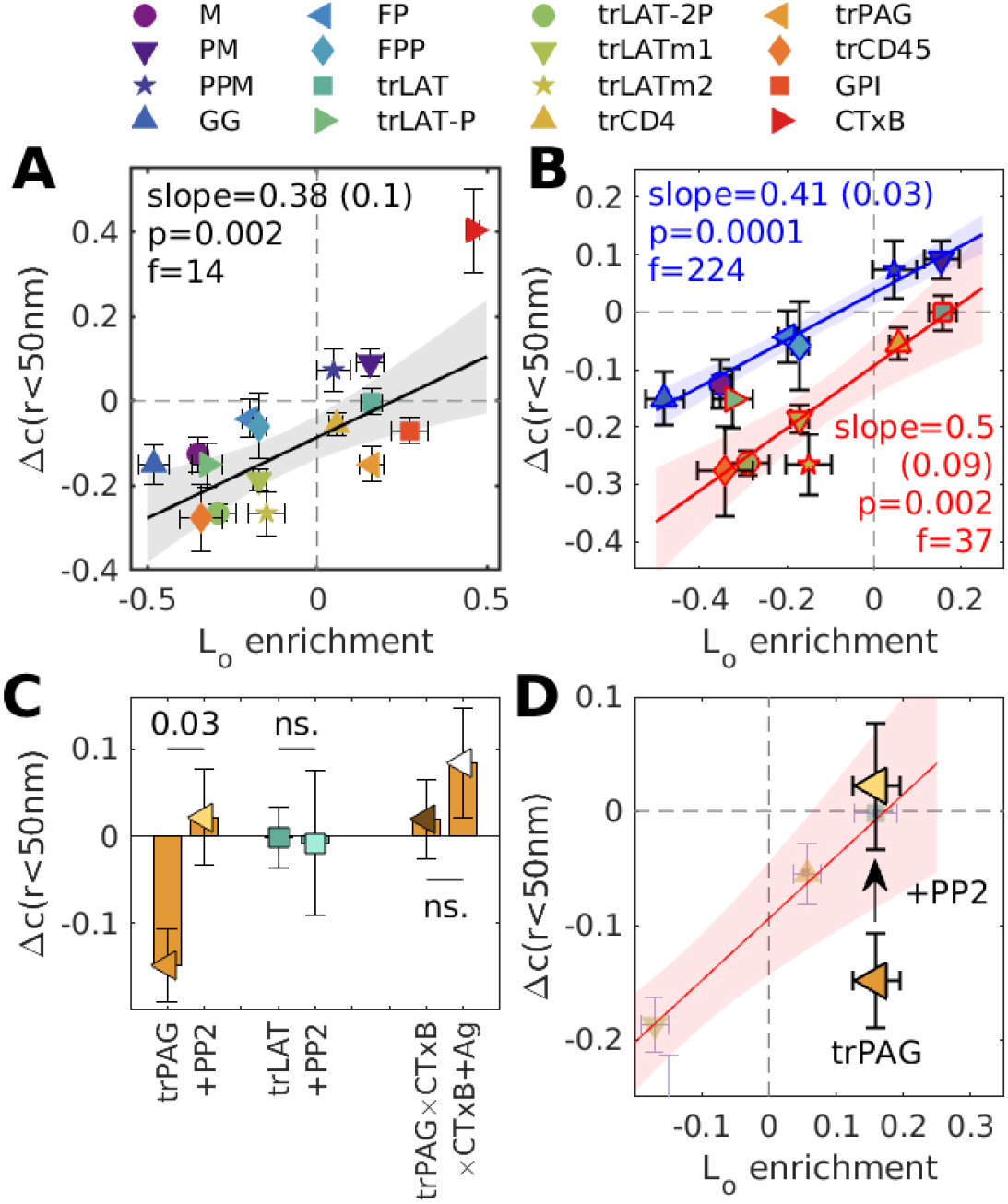
Anchor concentrations near receptor clusters are quantitatively predicted by GPMV measurements. (A) Changes in the magnitude of colocalization between anchors and BCR upon clustering in cells (Δc(r<50nm) from Fig 2B) plotted vs. Lo enrichment in isolated GPMVs (Fig 3B). (B) A subset of the data in A, separated by anchor class with peripheral anchors at the inner leaflet outlined in blue and transmembrane peptides outlined in red. Trends in A, B are fit to a linear model and the significance is assessed with a p-value vs. the null hypothesis of no correlation (lower values indicate greater significance), and an f-statistic that reports how well the variance in the data is described by the linear fit (higher values indicate greater significance). Shaded regions in A-B indicate the 95% confidence interval of the linear fit. (C) Changes in cross-correlation amplitudes (Δc(r<50nm)) for the specified probe-pairs both in the presence and absence of pre-treatment with PP2. trPAG was also imaged alongside clustered CTxB (×CTxB) in cells with (+Ag) and without BCR clustering. (D) PP2 treatment brings trPAG partitioning in-line with the correlations for transmembrane anchors redrawn from B.

Considering points corresponding to peripheral anchors and transmembrane anchors separately increases the significance of the linear model substantially (Fig 4B). Grouping points according to class also to brings experimental errors in line with the scatter of points around the linear trend, indicating that Lo phase partitioning in GPMVs is sufficient to explain the observed enrichment in BCR clusters (or vice versa) within anchor classes. The linear model describing inner leaflet peripheral anchors passes close to the point (0, 0), indicating that peripheral probes that partition equally between phases in GPMVs also show no preference for BCR clusters. The linear model describing transmembrane peptides is systematically shifted to lower values across the whole range of GPMV ordered phase affinities, indicating that transmembrane anchors experience a repulsive force from BCR clusters beyond that experienced by peripheral anchors. This possibly arises from steric hindrance of transmembrane domains within the crowded transmembrane environment at the center of these clusters. The slopes of these linear trends are close to 0.5, indicating that the magnitude of probe enrichment or depletion at BCR clusters is close to half that of Lo domains in GPMVs. We note that this value is likely an underestimate of contrast at the center of BCR clusters due to the finite localization precision and size of the BCR clusters themselves as described in Supplementary Figure S8.

Fig 4B omits the trPAG anchor whose preference for BCR clusters in cells represents a clear outlier from the linear trend. trPAG is markedly excluded from BCR clusters even though it partitions with Lo domains in isolated GPMVs to a similar degree as trLAT. trPAG is derived from PAG/CBP, an adaptor protein that plays important roles in downregulating BCR signaling (Kalland et al., 2012). To explore the role of signaling in trPAG localization, we clustered BCR in the presence of the Src kinase inhibitor PP2 and found trPAG to be enriched in BCR clusters, even though this treatment does not impact the distribution of trLAT (Fig 4C). We determined that the phase preference of trPAG is not directly impacted by BCR activation by monitoring its co-localization with clusters of CTxB in both stimulated and unstimulated cells (Fig 4C). Clustered CTxB has previously been shown to stabilize domains with Lo-like characteristics (Stone et al., 2017). Instead, BCR activation leads to the recruitment of trPAG to adhesive structures labeled by Focal Adhesion Kinase (Supplementary Figure S9). This finding, along with the PP2 dependence of trPAG co-localization with BCR leads us to conclude that protein-interactions sequester trPAG from signaling BCR clusters. Importantly, PP2 treatment brings Δc (r<50nm) for trPAG into agreement with the trend of Fig 4B (Fig 4D).

### Lo phase enrichment predicts relative anchor mobility

Fluorescent localizations used to reconstruct images and characterize BCR and anchor spatial distributions can also be used to quantify anchor mobility by measuring the temporal evolution of the anchor’s auto-correlation function, g(r, τ). g(r, τ) reports on the probability of finding pairs of localizations of the *same* molecule type separated by a distance r and a time-interval τ. This approach is similar to quantifying diffusion using single particle tracking (SPT) but has the advantage that segments are not explicitly linked to build trajectories, enabling unambiguous determination of mobility parameters when probes are imaged at high density. We find that g(r) at each τ is minimally described as a superposition of two states with distinct mean squared displacements (MSDs), and the evolution of these two states in τ reveals that one is highly confined and one is more diffusive over the range of time-intervals probed (15 ≤ τ ≤ 200ms). See Materials and Methods and Supplementary Figure S10 for methodological details, fits to g(r), MSD(τ) curves, and diffusion coefficients for several representative cells. A summary of all fit parameters extracted from probes imaged both before and after BCR crosslinking are supplied in Supplementary Figure S11. Overall, there is no clear trend between the Lo enrichment of anchors in GPMVs and mobility parameters describing the absolute diffusion coefficient of their mobile state (D) or the percentage of confined molecules (α) evaluated at the shortest time-interval (τ=15ms) (Figure 5A). Instead, anchor mobility differs mainly according to the position of the fluorescent tag (Supplementary Figure S12). These findings are in good general agreement with past work showing that membrane protein mobility is largely dictated by the size of the anchor, which is comparable across our pannel of anchors, and the presence of obstacles, which differs on the intra- and extracellular faces of the plasma membrane (A. Camley and H. Brown, 2013; Fujiwara et al., 2002; Guigas and Weiss, 2016; Kenworthy et al., 2004; Knight et al., 2010; Saffman and Delbrück, 1975). We next asked if BCR clustering and subsequent cellular activation alters anchor mobility, and if this effect correlates with Lo affinity. Figure 5B shows how the average diffusion coefficient varies with τ for GG and PPM, two anchors with similar absolute mobility but disparate partitioning behavior. For GG, BCR activation results in a small but systematic shift to larger diffusion coefficients (faster diffusion) across time-intervals, while diffusion coefficients are unchanged by cellular activation for PPM. The ratios of diffusion coefficients before and after BCR clustering for all anchors are summarized in Fig 5C, revealing a statistically significant correlation with Lo enrichment in GPMVs. BCR activation results in faster diffusion for anchors that partition to Ld domains (Lo enrichment<0), while diffusion is slower or remains unchanged for anchors that prefer Lo domains (Lo enrichment>0). We also find that confinement is reduced overall after BCR activation, with Lo-favoring anchors experiencing the greatest reduction in confinement (Supplemental Figure S13). This approach interrogates anchors imaged across the cell surface, with the vast majority of anchors localized away from BCR clusters. Thus, the direct physical impact of BCR clusters is likely minimal. Rather, it seems likely that stimulation-induced changes, such as in actin structure and cytoskeleton-membrane coupling, may contribute to the observed trend (Machta et al., 2011; Pore and Gupta, 2015). A similar trend was recently reported for Lo- and Ld-preferring anchors upon FcεRI stimulation in RBL-2H3 mast cells (Bag et al., 2021). A clear outlier to the trend of Fig 6B is CTxB. Past work suggests that diffusion of this anchor is dependent on membrane curvature (Kabbani et al., 2020), which is not likely to impact the mobility of other anchors due to their small size.

**Figure 5:**
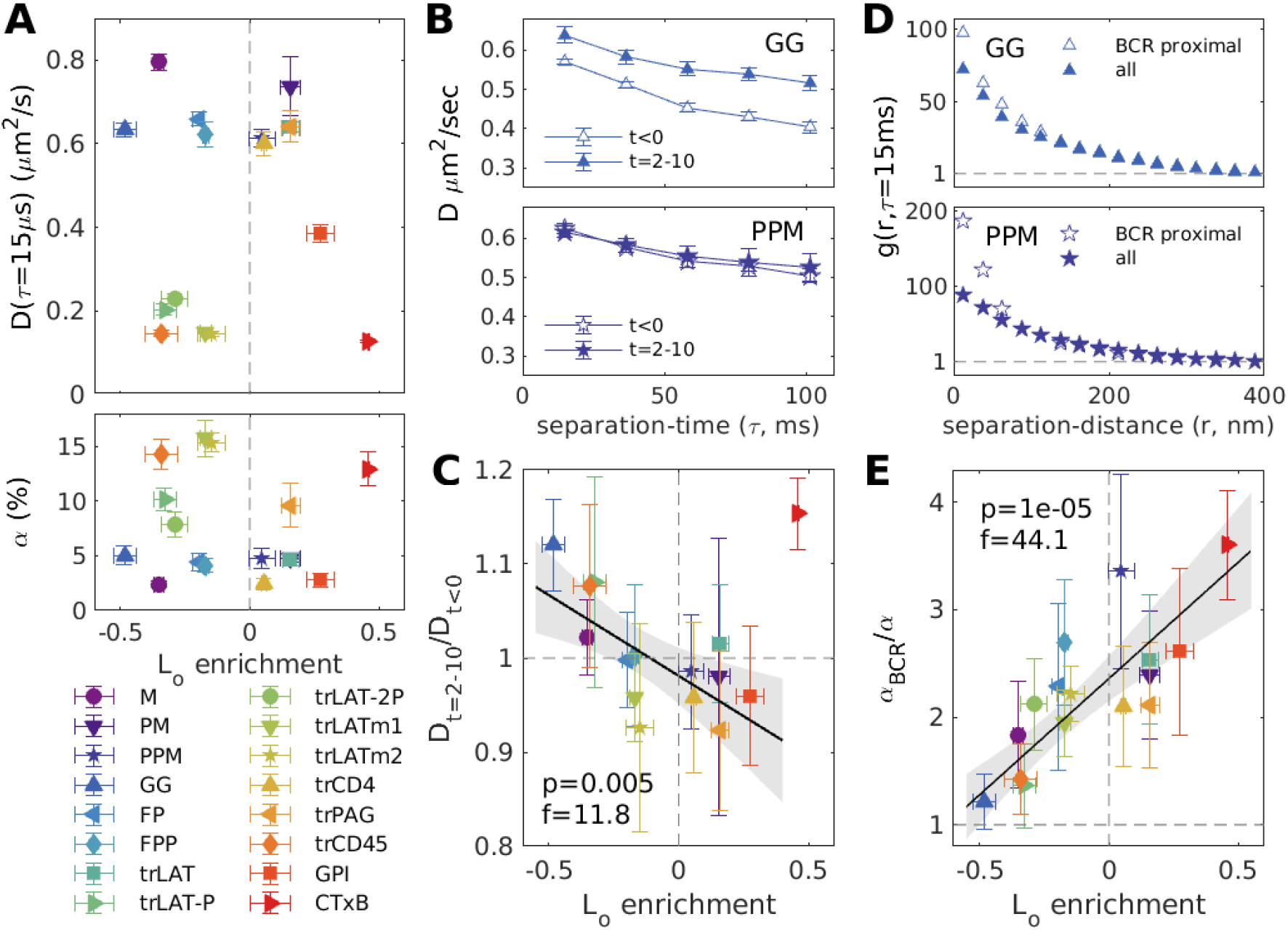
Relative anchor mobility is predicted by Lo enrichment. (A) Absolute diffusion coefficients (D) (top) and confined fractions (α) (bottom) for anchors observed 2-10 min after BCR clustering as determined from time-resolved auto-correlation functions (g(r, τ)) as described in Methods. (B) The diffusion coefficient (D) of the mobile fraction as a function of separation time τ from images acquired before (t<0min) and after (t=2-10min) BCR crosslinking. (C) Ratios of diffusion coefficients before and after BCR crosslinking for τ=15ms. The point corresponding to CTxB (red triangle) is excluded from the linear fit. A similar plot comparing α before and after crosslinking is presented in Supplementary Figure S13. (D) g(r, τ) functions showing the mobility of all anchors (all) or the subset of anchors localized within 100nm of a BCR cluster (BCR proximal) at τ=15ms. Curves are averages over all cells expressing the specified probe imaged 2-10min after BCR clustering. (D) Ratios of the confined fractions for all anchors (α) and BCR proximal anchors (α_BCR_) for τ=15ms. All points are included in the linear fit, but trLAT-2P is excluded due to poor statistics in unstimulated cells. A similar plot comparing D overall to D proximal to BCR is in Supplementary Figure S13. Trends in B, D are fit to a linear model and the significance is assessed with a p-value and an f-statistic as described in Fig 4. Shaded regions in B, D indicate the 95% confidence interval.

**Figure 6:**
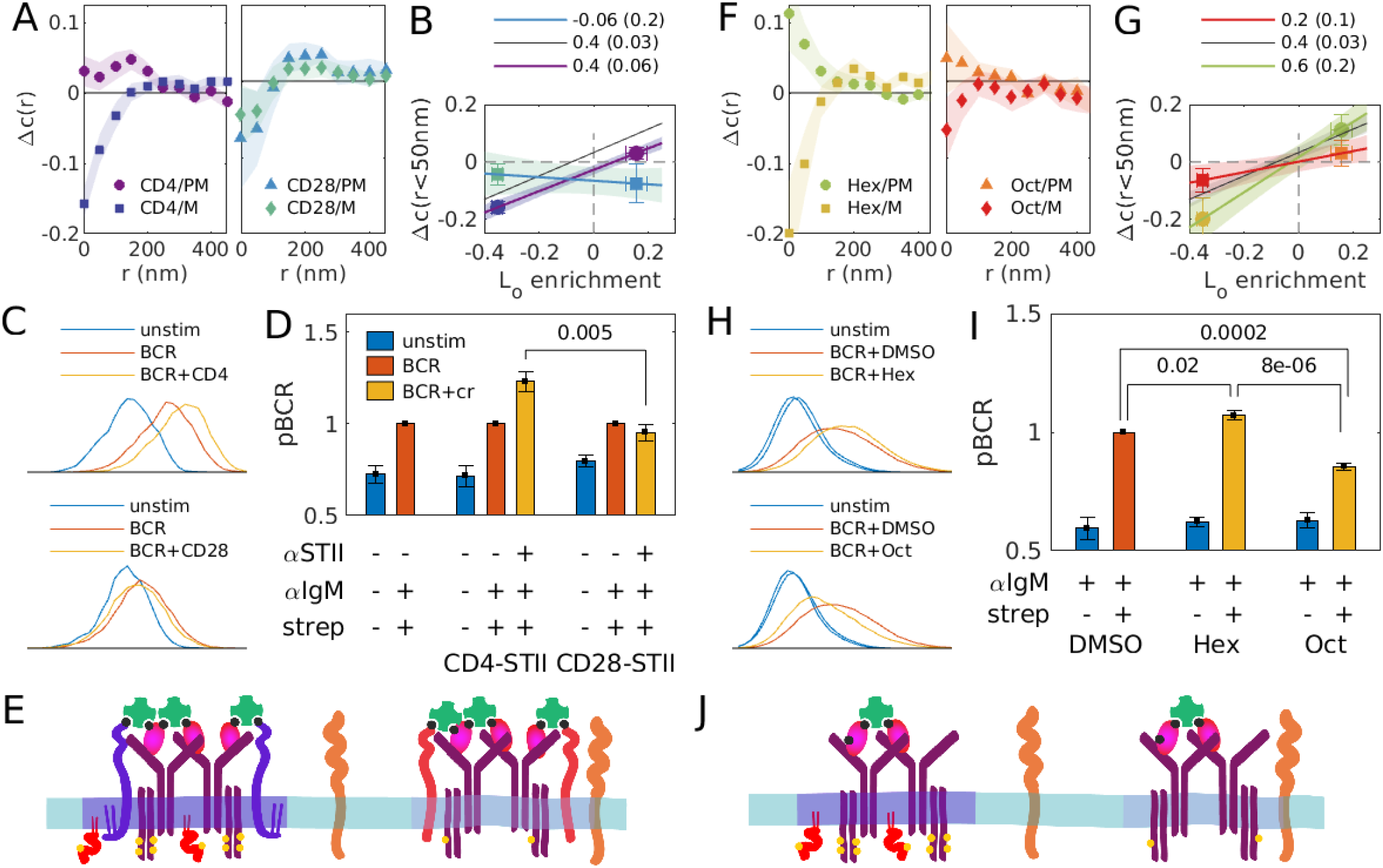
BCR phosphorylation is modulated by perturbations of domain contrast. (A) Average change in cross-correlation functions (Δc(r)) between BCR and either PM or M anchors, before and after BCR is co-ligated with minimal co-receptors trCD4cr (left) or trCD28cr (right). Post stimulation cross-correlation functions are tabulated from localizations acquired between 2 and 10 min after co-ligation with minimal co-receptors. Error bounds indicated by the shaded region represent the SEM over cells. (B) Co-ligation induced change in M or PM probe sorting with respect to BCR (Δc(r<50nm) from Fig 6A) plotted vs. Lo enrichment of M or PM in isolated GPMVs (Fig 3B). The slope of the line connecting these points reports the extent to which sorting in cells quantitatively compares to sorting in phase separated vesicles. Here, shaded regions are tabulated by propagating errors from measured points. The black line is the linear fit for all inner leaflet anchors reproduced from Fig 4B. (C) Representative histograms from flow cytometry measurements of pBCR staining over a population of cells expressing trCD4cr (top) or trCD28cr (bottom), where suspension cells were fixed and stained before (unstim) or after either BCR clustering (BCR) or co-ligation of BCR and minimal co-receptors as described in Methods. (D) Summary of pBCR levels of various treatments normalized to the BCR cross-linking condition for untransduced, trCD4cr-expressing, and trCD28cr-expressing cells. (E) Schematic summary of experimental findings and proposed model: BCR clusters with higher domain contrast also support higher levels of BCR tyrosine phosphorylation, possibly due to sorting of regulatory kinases and phosphatases. (F-J) Results of imaging (F,G) and flow cytometry (H,I) experiments conducted in parallel with experiments described in A-E, where here instead of co-ligation with co-receptors, untransduced cells were pre-treated with either hexadecanol or octanol before BCR crosslinking as described in

We next compared the mobility of all anchors in stimulated cells to the subset of anchors that pass within 100nm of a BCR cluster to isolate effects on diffusion that may arise from the distinct membrane regions around clustered receptors. Fig 5D shows how correlation curves for BCR-proximal probes compare to curves generated from all probe localizations at τ=15ms for GG and PPM. In both cases, the most notable effect is that BCR-proximal anchors have increased auto-correlation amplitudes at short r, indicating that anchors are more likely to travel short distances when they are near BCR clusters. Fitting correlation functions for BCR-proximal anchors indicates that this change can be attributed to a larger population of confined anchor molecules close to BCR clusters (α_BCR_) than are present in the membrane overall (α) without meaningful changes in the diffusion coefficient of the mobile component (Supplementary Figure S13). A similar effect is observed across all anchors, although the relative magnitude of the confined population (α_BCR_/α) varies. A highly significant trend is observed when this ratio is considered alongside Lo enrichment measured in GPMVs (Fig 5E). Specifically, the most Lo-enriched anchors are 3-4 times more likely to be confined near a BCR cluster than in the membrane overall, while strongly Ld-enriched anchors are minimally impacted by the presence of a BCR cluster. This trend holds despite the lack of a significant correlation between Lo partitioning and either α or α_BCR_ when considered individually (Supplementary Figure S11). This striking relationship suggests that stabilization of an ordered membrane environment around clustered BCR can control not only recruitment but also the retention of relevant membrane components.

### Modulating domain partitioning impacts BCR function

BCR is phosphorylated at cytoplasmic tyrosine residues by kinases such as Lyn and Syk and dephosphorylated by phosphatases such as CD45 and SHP-1. Some of these regulatory proteins are anchored to the membrane via motifs that are sorted by BCR clusters according to their Lo preference (Fig 4B). This suggests that the ability of regulatory proteins to phosphorylate or dephosphorylate BCR is dictated in part through their access to BCR clusters based on their partitioning into membrane phases. To test this hypothesis, we identified perturbations that altered the extent of phase-mediated sorting at BCR clusters and monitored the impact on BCR tyrosine phosphorylation levels (“pBCR”), as summarized in Figure 6.

Our first strategy to modulate the magnitude of phase-mediated sorting at BCR clusters is to incorporate additional components into BCR clusters that either favor or oppose formation of Lo membrane domains. To accomplish this, BCR was co-ligated with one of two minimal engineered “co-receptors” transduced to express at the plasma membrane of CH27 cells. One minimal co-receptor has the palmitoylated transmembrane domain of CD4 which incorporates into the Lo phase in GPMVs (trCD4cr, Fig 3), while the second has the transmembrane domain of CD28, which lacks palmitoylation sites and is excluded from the Lo phase in GPMVs (trCD28cr, Supplementary Figure S14). Both minimal co-receptors lack intracellular signaling domains and contain an extracellular Strep-tagII (STII) used to co-ligate with IgM BCR using biotinylated antibodies against STII and IgMμ respectively, as described in Methods. Fluorophores decorating fAb αIgMμ and the transduced co-receptors assemble into tight puncta upon addition of streptavidin (Supplementary Figure S15). The partitioning of components with respect to these puncta was assayed using PM and M, anchors that sort strongly with the Lo and Ld phase, respectively. Quantifying cross-correlations between BCR and these anchors shows that BCR clusters co-ligated with trCD4cr more effectively sort PM and M than BCR clusters that incorporate trCD28cr (Fig 6A-B), indicating that co-receptors impact the contrast of phase-like domains. BCR tyrosine phosphorylation (pBCR) was assayed by flow cytometry of suspended cells where BCR was co-ligated with minimal co-receptors. pBCR intensity is enhanced in cells expressing trCD4cr as compared to cells expressing trCD28cr (Fig 6C-D). This result supports the previous assertion that some endogenous BCR co-receptors function, at least in part, by altering the membrane properties at BCR clusters (Cherukuri et al., 2001, 2004).

A second strategy to modulate phase-mediated sorting with respect to BCR clusters is to perturb membrane composition overall to either enhance or suppress membrane phase separation. To accomplish this, we incubated cells with either octanol or hexadecanol, hydrophobic compounds previously shown to lower or raise miscibility transition temperatures in isolated GPMVs (Machta et al., 2016). Cross-correlations between BCR and M or PM show that hexadecanol treatment results in BCR clusters that more effectively sort PM and M anchors, while octanol treatment results in BCR clusters that only weakly sort these anchors (Fig 6F-G), indicating that these treatments impact the contrast of phase-like domains. The specific treatments used have an equal and opposite impact on membrane phase transition temperatures in isolated GPMVs and also have equal magnitude, opposing effects on BCR domain contrast. Flow cytometric analysis indicates that hexadecanol treatment enhances and octanol treatment inhibits phosphorylation of clustered BCR (Fig 6H-I).

Overall, we modulated the strength of phase-mediated sorting at BCR clusters in two ways: extrinsically by altering the protein composition of BCR clusters, and intrinsically by shifting the membrane phase transition temperature. In both cases, modulations that increase the contrast of domains also lead to higher levels of BCR phosphorylation while modulations that decrease contrast lead to lower levels of BCR phosphorylation, as illustrated schematically in Fig 6E and J.

## DISCUSSION

Clustering the BCR leads to assembly of a signaling platform with a distinct protein and lipid composition, establishing a local environment that facilitates functional interactions between signaling proteins. These signaling platforms are held together by signaling and scaffolding proteins, but also exhibit distinct local membrane environments that influence downstream signaling responses. While a large number of past studies link BCR activation and downstream signaling to biochemically defined “lipid rafts”, the lack of a structural correlate or mechanistic basis in intact cells impedes integrating these observations into a predictive model of the immune response.

Here we present a systematic comparison of anchor organization in parallel but independent experiments in cells and vesicles that establishes phase separation as an underlying mechanism for assembly of functional domains. We show that membrane-mediated forces are sufficient to alter composition proximal to BCR, with membrane-anchored probes recruited or excluded from these signaling platforms despite the absence of any specific protein interaction domains (Fig 1). A role for phase separation naturally emerges because a diversity of membrane anchoring motifs are examined and because quantitative methods can directly measure protein concentration in both systems, the parameter that is determined by the thermodynamics of this phase transition.

We observe a remarkable convergence between observations of probe enrichment in BCR clusters (Fig 2) and probe enrichment in Lo domains in isolated GPMVs (Fig 3), validating past implications (Gold and Reth, 2019; Gupta and DeFranco, 2007; Pierce and Liu, 2010; Sohn et al., 2006; Stone et al., 2017) that BCR signaling platforms are extended ordered domains (Fig 4). Moreover, the phase preference also predicts aspects of probe mobility (Fig 5), in particular the retention of probes at BCR clusters. Together these findings support the conclusion that liquid-liquid phase separation underlies lipid raft heterogeneity within B cell membranes.

### A new perspective on ordered domains in cell membranes

Our observations, combined with other recent studies (Kinoshita et al., 2017; Koyama-Honda et al., 2020; Li et al., 2020; Stone et al., 2017), put to rest a long history of debate regarding the existence of ordered membrane domains often referred to as ‘lipid rafts’ in cells. Our findings clearly demonstrate that ordered domains can and do exist in intact cells, that they can emerge in response to stimuli (Fig 1), and that their presence impacts the concentration and dynamics of membrane proteins (Figs 4, 5). Although macroscopic separation of Lo and Ld phases does not occur in intact plasma membranes, insights from model membranes explain how phase separation underlies emergent structure in living systems.

Nanoscale heterogeneity has been consistently reported in model membranes that can be driven to macroscopic phase separation through modest changes in temperature or composition (Bloom and Thewalt, 1995; Heberle et al., 2010; Ingólfsson et al., 2014; Oldfield and Chapman, 1972; Pathak and London, 2015). Past studies (Ge et al., 2003; Veatch et al., 2008), including several recent contributions (Heberle et al., 2020; Li et al., 2020; Schneider et al., 2017), demonstrate that nanoscopic structure also exists in isolated plasma membranes, even where no macroscopic phase separation can be observed. Past work also indicates that domains can be extended from the nano to the micron scale through external forces acting on membranes. For example, lipid probes that prefer Lo or Ld phases can be sorted by regions of high curvature in single phase membranes close to a miscibility phase transition even in the absence of proteins, and this effect is amplified when proteins are included that couple to both curvature and membrane phases (Heinrich et al., 2010; Sorre et al., 2009). Extended phase-selective domains also form when proteins or lipids are clustered through adhesion (Zhao et al., 2013) or when membranes are coupled to protein complexes (Liu and Fletcher, 2006; Manley et al., 2008), cytoskeletal scaffolds (Arumugam et al., 2015; Honigmann et al., 2014; Vogel et al., 2017), or phase separated protein droplets (Chung et al., 2021; Lee et al., 2019). While the detailed physical mechanisms underlying these phenomena remain an area of active debate (Schmid, 2017), it is clear that single phase model membranes near macroscopic Lo-Ld phase transitions are highly susceptible to perturbations that organize components with specific phase preferences. Our results reinforce that the same principle extends to living cell membranes.

While membrane phase separation contributes to the composition of BCR clusters, these domains are not held together solely by phase-mediated interactions (Gauld et al., 2002; Mattila et al., 2016; Packard and Cambier, 2013; Tolar et al., 2009). As such, they exhibit properties that differ from coexisting phases in isolated vesicles. In our results, this difference manifests in reduced magnitude of enrichment/depletion of anchors in BCR clusters compared to phase separated domains within vesicles (Fig 4). Similar effects are evident in model membranes with extended stabilized domains, where the contrast of phase-marking probes is reduced when the membrane is further from its macroscopic miscibility phase transition (Sorre et al., 2009; Zhao et al., 2013). The reduced contrast between enrichment in Lo phases of isolated plasma membrane vesicles and BCR clusters is roughly 2-fold (slopes are 0.4 and 0.6 in Fig 4B).

We propose that the marginal loss of contrast inherent to tuning away from macroscopic phase separation is functionally outweighed by the adaptability the membrane acquires in this unique physical state. Near, but not at, phase separation, extended domains similar to phases can assemble or disassemble in response to external stimuli rather than thermodynamic parameters of the membrane itself (Figure 7). Protein assemblies coupled to membranes localize and extend existing nanoscale structure, establishing functional compartments that are stable in both space and time. Here we demonstrate concrete examples whereby domain contrast is responsive to the identity of scaffolding proteins in the form of minimal co-receptors (Fig 6). Despite this sensitivity to the stimulus, thermodynamic parameters of the membrane still play an important role by tuning the magnitude of membrane phase-mediated interactions and changing the composition of domains (Machta et al., 2012). In isolated vesicles, numerous factors impact the location of the macroscopic phase boundary, including cell type, perturbations to lipid homeostasis, incorporation of drugs, and the presence or loss of leaflet asymmetry (Cammarota et al., 2020; Levental et al., 2017, 2020b; Machta et al., 2016; St. Clair et al., 2020). In principle, these factors may all be used by cells to tune the magnitude of phase-mediated enrichment during cellular processes. In the current work, we used a calibrated perturbation of membrane transition temperature and demonstrated a corresponding change in domain contrast (Fig 6). Bringing these ideas together, membrane domains enable a tunable response by integrating the thermodynamic state of the membrane and the magnitude of the applied stimulus.

**Figure 7:**
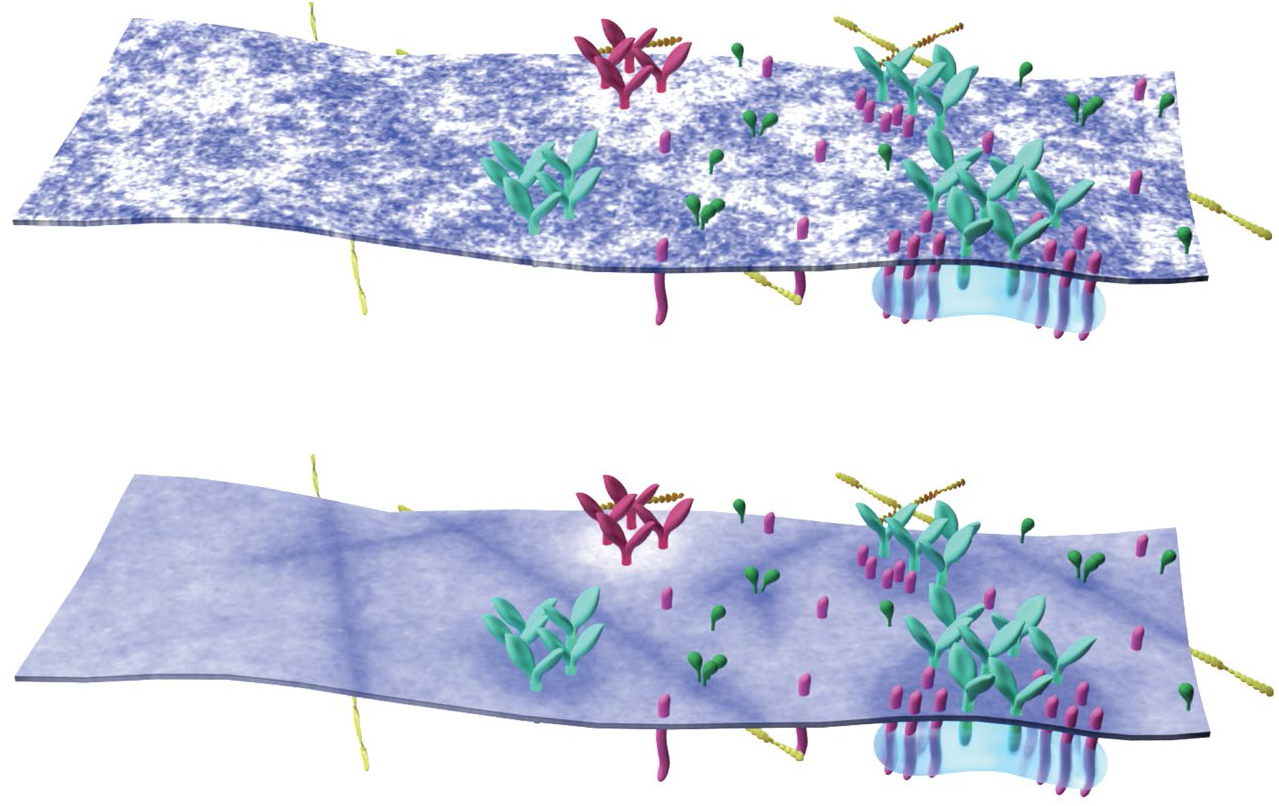
A model for adaptive membrane organization. Nanoscale membrane domains preferentially assemble at protein scaffolds to establish functional compartments in the plasma membrane. (top) Instantaneous snapshot of a heterogeneous membrane containing protein structures of increasing complexity, including cortical actin, clustered receptors, adaptor proteins, and membrane proximal signaling assemblies. This membrane contains numerous domains of different sizes and shapes, some of which surround protein rich structures. (bottom) Same membrane drawn above but now membrane composition is averaged over time to show continuous areas of altered local concentration, demonstrating how nano-scale structure can give rise to distinct local membrane environments and define membrane compartments. This is a representation of one specific physical model of membrane heterogeneity (Supplementary Fig S16) but we expect this effect to be general across models.

### Stabilized domains in signaling function

In past work we found that receptor kinases and their minimal anchors are enriched in BCR clusters, whereas phosphatases and their minimal anchors are excluded (Stone et al., 2017). This effect offsets the balance of kinase-phosphatase activity to favor receptor tyrosine phosphorylation. Here we show that the physical basis of sorting is phase separation (Fig 4) and that receptor phosphorylation is tuned by the extent of phase-mediated sorting (Fig 6).

These results allow us a context to apply a mechanistic interpretation to extensive past studies centered around how lipid rafts tune BCR signaling through organization of membrane components. For example, past work shows that the co-receptor CD19 enhances signaling when co-clustered with BCR, in part through modulating the composition and longevity of biochemically defined rafts (Cherukuri et al., 2001). Dynamic palmitoylation of the tetraspanin CD81 associated with CD19 was necessary for the enhancement of the BCR activation signal (Cherukuri et al., 2004). In the context of our current results, we can ascribe a specific mechanism to this effect as modulating the effective partitioning of BCR signaling complexes, changing the character of the associated membrane compartment and therefore interactions with downstream signaling mediators. We demonstrate this effect for the specific example of minimal co-receptors that result in altered anchor partitioning and BCR phosphorylation even in the absence of additional signaling domains. Factors that change the local composition of BCR signaling compartments, such as global changes in membrane composition that accompany B cell development and maturation (Karnell et al., 2005; Sproul et al., 2000), would be predicted to have similar effects. This is demonstrated here for the specific example of a calibrated perturbation of membrane transition temperature. Importantly, considering results within the rigorous framework of phase separation equips researchers with tools to stitch together findings isolated by specific cellular or stimulation contexts into a unified picture.

Changes in mobility imposed by domains may be another factor that influences signaling interactions. In many cases, the kinetics of interactions between signaling partners set the threshold for cellular activation in response to stimuli, where productive events require sustained assembly of a signaling complex that increases interaction dwell times. This concept is invoked by kinetic proofreading and kinetic segregation models of T cell receptor activation (Davis and van der Merwe, 2006; Ganti et al., 2020; McKeithan, 1995; Tischer and Weiner, 2019; Yousefi et al., 2019). Partitioning-dependent confinement of membrane anchors (Fig 5 D-E) increases their dwell time within domains, potentially increasing the probability of serial engagement and biasing signaling pathways towards activation. Co-confinement of signaling proteins within domains could be a general mechanism by which receptor clustering enhances signaling interactions (Koyama-Honda et al., 2020; Stone et al., 2017).

As a general factor influencing membrane organization, phase separation likely has a broad role in spatial regulation of membrane signaling pathways beyond those of clustered receptors. Lo partitioning is a direct consequence of anchor protein structure and posttranslational modification (Levental et al., 2010; Lorent et al., 2017), providing a possible explanation for why protein family members with similar catalytic domains but different membrane anchoring motifs accomplish different cellular functions. Through their particular anchors, membrane proteins can gain preferential access to different membrane compartments defined by the local membrane environment. Mutual affinity for membrane compartments effectively increases interactions between individual protein molecules, impacting their biochemistry and ultimately their functions. Because phase separation is equally able to enrich order- and disorder-preferring membrane components, we anticipate that both ordered and disordered compartments can regulate productive signaling.

The ordered domain established by BCR clusters is likely just one example of domains stabilized by signaling complexes in the plasma membrane. Indeed, any structure that contacts the membrane can in principle couple to a specific membrane compartment, presenting intriguing possibilities about how extended structures such as the actin cytoskeleton can broadly regulate membrane signaling pathways via interactions with membrane microstructure. For example, membrane-cytoskeleton contact sites that preferentially associate with a particular membrane compartment would influence the organization and dynamics of essentially all proteins throughout the membrane plane, based on their phase preferences (as sketched in Fig 7) (Machta et al., 2011). Changes in network structure, i.e. actin dynamics or ERM protein engagement, would produce global changes in membrane organization, impacting interactions across the cell surface (Shelby et al., 2016). Such changes to actin networks occur during signaling through the BCR (Pore and Gupta, 2015), and it seems likely that the changes observed in overall anchor mobility after BCR crosslinking (Fig 5B-C and Supplementary Figure S13) are a result of changes in actin coupling. While past work focuses primarily on the role of actin in restricting the mobility of specific proteins (Gasparrini et al., 2016), it seems possible that global changes in membrane domains could also play functional roles. Global changes in membrane domains are also expected when thermodynamic parameters within the membrane are altered, as could happen in response to slow changes in lipid metabolism or abrupt changes that occur during the cellular signaling response such as the loss of lipid asymmetry (Elliott et al., 2006) or the enzymatic conversion between lipid species (Marshall et al., 2000).

To conclude, we propose a new perspective on membrane domains in cellular processes. This view presents a refinement of the traditional picture of rafts as stable, pre-existing domains that can grow or coalesce in response to stimuli. Instead, we envision membrane phases acting as an adaptive solvent that is highly susceptible to forces acting in and on membranes. This solvent impacts the local concentration and dynamics of proteins within domains, shifting the balance of biochemical reactions and therefore membrane-mediated functions. These domains are sufficient to sort membrane components based on their phase preference even when components lack direct interactions with clustered proteins, and this sorting is sensitive both to external cues and to thermodynamic parameters within the membrane. In this way, the plasma membrane phase transition works in concert with other organizing principles to facilitate and modulate cell function.

## ACKNOWLEDGEMENTS

We thank Ben Machta, Thomas Shaw, Isabella Graf, Nirmalya Bag, and Barbara Baird for helpful conversations. Research was supported by grants from the NIH (GM110052 to SLV, GM134949 and GM124072 to IL), the NSF (MCB1552439 to SLV), the American Cancer Society (PF1800401CCE to SAS), the Volkswagen Foundation (grant 93091 to IL), and the Human Frontiers Science Program (RGP0059/2019 to IL).

## AUTHOR CONTRIBUTIONS

Conceptualization, SAS and SLV; Methodology, SAS,IL, SLV; Investigation, SAS and ICS, Formal Analysis, SAS and SLV, Resources KW and IL, Writing – original draft, SAS and SLV; Writing – review and editing, SAS, IL, SLV; Visualization, SAS and SLV; Supervision, IL and SLV;

## DECLARATION OF INTERESTS

The authors declare no competing interests.

## METHODS

### Cells

CH27 B-cells (mouse, Millipore Cat# SCC115, RRID:CVCL_7178), a lymphoma-derived cell line (Haughton et al., 1986) were acquired from Neetu Gupta (Cleveland Clinic). CH27 Cells were maintained in culture as previously described (Stone and Veatch, 2015). RBL-2H3 (rat, ATCC CRL-2256; RRID: CVCL_0591) and HEK-293 (HEK) (human, ATCC CRL-1573; RRID: CVCL_0045) cells were purchased from ATCC and cultured in medium containing 89% Eagle’s Minimum Essential Medium (EMEM), 10% FCS, and 1% penicillin/streptomycin at 37 °C in humidified 5% CO2.

### Probes and Conjugation

#### Constructs

The majority of anchor constructs used in this study were derived from existing anchor-fluorescent protein fusion constructs, with the fluorescent protein sequence replaced by that of mEos 3.2 via standard molecular cloning. All constructs were expressed through transfection of mammalian expression plasmids. The original plasmids were obtained from a variety of sources, and detailed sequence and source information for all anchor constructs is provided in Supplementary Table 1. FAK-eGFP plasmid DNA was obtained from the Baird-Holowka lab.

Constructs for minimal “co-receptors” trCD4cr and trCD28cr used for experiments described in Fig 6A-E were derived from truncated, fluorescently tagged versions of chimeric antigen receptor constructs that have been described previously (Liu et al., 2016). Sequences consist of a CD19-specific scFV followed by the 9 aa Strep-tagII sequence (NWSHPQFEK) fused with a Gly/Ser (G4S)2 linker, a short 12 aa IgG4 hinge sequence, a transmembrane domain followed by a short cytoplasmic overhang sequence from either CD4 (M397-Q428 of human CD4) or CD28 (F153-D190 of human CD28), and an intracellular eGFP tag. Constructs were cloned in the lentiviral epHIV7 vector (Yam et al., 2002). Lentiviral supernatants were produced in HEK 293T cells transfected with lentiviral vector plasmids along with PAX2 and VSVG packaging plasmids. Viral particles were concentrated via ultracentrifugation before use for transduction.

#### Lipid dyes

FAST-DiO, FAST-DiI and DiD were purchased from ThermoFisher.

#### Antibodies, protein reagents, and dyes

Goat anti-Mouse IgM(μ) Fab fragments were obtained from Jackson Immunoresearch (Jackson ImmunoResearch Labs Cat# 115-007-020, RRID:AB_2338477) and were conjugated with biotin and silicon rhodamine (SiR) NHS esters (Invitrogen: B1582, Spirochrome: SC003) to produce “SiR-biotin-Fab”. Non-fluorescent biotin-conjugated Goat anti-Mouse IgM(μ) Fab fragments were also obtained from Jackson Immunoresearch (Jackson ImmunoResearch Labs Cat# 115-067-020, RRID:AB_2338587). Anti-GFP nanobody used to label FAK-eGFP was from Chromotek (Chromotek: gt-250) and was conjugated to JaneliaFluor 646 through incubation with JF646 NHS ester (gift of Luke Lavis, Janelia Farms). Cholera Toxin subunit B (CTxB) was from Sigma (Sigma: C9903) and was conjugated with Cy3B NHS ester (Cytiva: PA63101) or with both SiR and biotin as with SiR-biotin-Fab. Rabbit monoclonal antibody against phospho-CD79A (Tyr182) was purchased from Cell Signaling (Cell Signaling Technology Cat# 5173, RRID: AB_10694763). Biotin-labeled mouse monoclonal antibody against the Strep-tagII (“NWSHPQFEK”) was purchased from Genscript (GenScript Cat# A01737, RRID:AB_2622222). Alexa Fluor 568 streptavidin was from ThermoFisher (Thermo Fisher Scientific Cat# S-11226, RRID:AB_2315774). Alexa Fluor 647 goat anti-rabbit IgG and Fcγ-specific goat anti-human IgG secondary antibodies were purchased from Jackson Immunoresearch (Jackson ImmunoResearch Labs Cat# 111-005-144, RRID:AB_2337919, Cat# 109-005-170, RRID:AB_2810885). Recombinant VCAM-1/human Fc chimera protein was purchased from R&D Systems (R&D Systems Cat# 862-VC-100).

#### Dye conjugation of labeling proteins

Antibodies, Fab fragments, and CTxB were all labeled via NHS ester chemistry using the same basic procedure. Amine-reactive dye succinimidyl esters were added to labeling protein solutions (.5 - 2 mg/ml) in 2 - 8x molar excess from stock solutions in DMSO. Reactions were buffered by 0.01M NaH2PO4 with 0.01M NaH2CO3 at pH 8.5 and proceeded for 1 hr at room temperature with rotation. After incubation, the modified proteins were purified and separated from unbound dye through a gel filtration column (GE Healthcare illustra NAP Columns from Fisher: 45-000-151) in PBS with 1mM-EDTA. The modified proteins were further purified by centrifugation in a Vivaspin-500 Polyethersulfone concentration spin column (Vivaproducts: VS0191, VS0121, VS0131, VS0141) with an appropriate molecular weight cutoff (3k-100k) of approximately half the protein molecular weight or lower. The degree of labeling was estimated using the absorbance spectrum of the conjugated protein.

SiR-biotin-Fab and SiR-biotin-CTxB were conjugated to SiR and biotin using a two-step procedure, where first unlabeled proteins were reacted with a mixture of SiR and biotin NHS esters in 5x and 12x molar excess, respectively, and purified as described above. A second round of labeling with SiR NHS ester alone was then performed, with 10x excess dye for SiR-biotin-Fab and 4x excess for SiR-biotin-CTxB. The degree of labeling for these conjugates was 2.5 SiR per protein for SiR-biotin-Fab and 2.1 SiR per protein for SiR-biotin-CTxB.

### Preparation of CH27 cell samples for super-resolution imaging

#### Transient transfection and viral transduction

CH27 cells were transiently transfected with plasmid DNA encoding membrane anchor constructs by Lonza Nucleofector electroporation (Lonza, Basel, Switzerland) with electroporation program CA-137. Typically, 10^6^ CH27 cells were transfected with 1 μg of plasmid DNA. Cells were allowed to recover in growth media in the incubator for 18-24 hours before preparation for imaging.

Minimal co-receptors trCD4cr and trCD28cr were expressed in CH27 cells via viral transduction for the experiments described in Fig 6A-E. Production of viral particles is described above. CH27 cells were transduced with concentrated viral supernatants at a 1:1 ratio of supernatant to cell growth media in a 6-well plate with 10^6 cells per well. To maximize transduction efficiency, polybrene (Sigma: TR-1003) was added to the media as a linker molecule at a final concentration of 8 μg/ml and the plate was spun for 2500 rpm for 30 min prior to incubation for 24 hours under cell culture conditions (37°C, 5% CO_2_). Media was exchanged, followed by another 48 hour incubation to allow cells to express the construct. Transduction was assessed based on brightness and membrane localization of the co-receptor eGFP tag via fluorescence microscopy. Transduced cells were then maintained in cell culture as normal. Transient transfection of M and PM anchors was performed on stably transduced trCD4cr or trCD28cr-expressing cells for the imaging experiments described in Fig 6A-B.

#### Live cell sample preparation

In most cases, transiently transfected cells expressing anchor constructs were transferred directly to glass-bottom Mattek culture dishes to recover overnight for next-day sample preparation. In some cases, cells recovered overnight in flasks and were plated on Mattek dishes on the day of sample preparation. BCR was labeled by incubation with 5 μg/ml SiR-biotin-Fab in culture medium for 10 min at room temperature. For live cell imaging experiments, BCR labeling was performed immediately prior to imaging.

When Cy3B-CTxB was imaged alongside SiR-biotin-Fab-labeled BCR instead of a transiently-expressed anchor construct, Cy3b-CTxB was added to the SiR-biotin-Fab labeling solution at a concentration of 5 μg/ml. When SiR-biotin-CTxB was imaged in conjunction with trPAG, SiR-biotin-CTxB was added at 5μg/ml instead of SiR-biotin-Fab in culture medium for 10 min prior to imaging.

For imaging experiments involving treatment with alcohols only (Fig 6F-G), cells transfected with anchor constructs were grown overnight in flasks and plated on Mattek dishes coated with VCAM-1 for 10-20 min prior to imaging. Plating cells on VCAM-1 coated dishes was performed to promote cell adhesion and maintain membrane flatness, which can be compromised by exposure to alcohols and/or vehicle DMSO. BCR was then labeled with SiR-biotin-Fab as described above. VCAM-1 coated dishes were prepared by incubation of oxygen plasma cleaned Mattek dishes with Fcγ-specific goat anti-human IgG antibody in PBS at 100 μg/mL for 30 min at room temperature. Dishes were blocked for 30 min with a 5% BSA solution, then incubated for 1 hour at room temperature with recombinant VCAM-1/human Fc chimera protein at 10 μg/mL, and rinsed thoroughly before use.

For imaging experiments on trCD4cr and trCD28cr transduced cells (Fig 6A-B), cells were transiently transfected with anchor constructs and plated on Mattek dishes. BCR and co-receptors were labeled in a single incubation step with a mixture of 1 μg/ml SiR-biotin-Fab and 5 μg/ml biotinylated mouse anti-Strep-tagII mAb for 10 min at room temperature immediately prior to imaging.

#### Fixed cell sample preparation

For fixed cell imaging experiments, BCR was labeled with biotin-Fab using the same procedure as live cell imaging experiments. Cells were then rinsed with HEPES-buffered salt solution with BSA (HBSS-BSA: 30mM HEPES, 5.6 mM glucose, 100 mM NaCl, 5 mM KCl, 1 mM KCl, 1 mM MgCl2, 1.8 mM CaCl2, .1% w/v BSA, pH 7.4) and were either chemically fixed or BCR was clustered through addition of 1 μg/ml streptavidin for 6 min. Stimulated and unstimulated samples were rinsed with PBS, then fixed (2% PFA .15% glutaraldehyde in PBS) for 10 min at room temperature. Samples were blocked and permeabilized using block solution (PBS with .5% w/v BSA, .05% v/v fish gelatin, .01% Triton-X 100) for 1 hour. For FAK labeling, cells were transiently transfected with FAK-eGFP plasmid DNA and were labeled with JF646 anti-GFP nanobody (1 μg/ml) for 30 min at room temperature after fixation, permeabilization, and blocking. Samples were then thoroughly rinsed with block solution and imaged.

### Single molecule imaging of B cells

Imaging was performed using an Olympus IX81-XDC inverted microscope. TIRF laser angles where achieved using a cellTIRF module, a 100X UAPO TIRF objective (NA = 1.49), and active Z-drift correction (ZDC) (Olympus America) as described previously (Stone and Veatch, 2015; Stone et al., 2017). SiR and JF646 were excited using a 647 nm solid state laser (OBIS, 100 mW, Coherent) and Cy3B was excited using a with a 561 nm solid state laser (Sapphire 561 LP or OBIS 561-120FP, Coherent). Photoactivation of mEos3.2 was accomplished with a 405 nm diode laser (CUBE 405-50FP, Coherent) and simultaneously excited with the 561 nm solid state laser. Simultaneous imaging of SiR or JF646 and mEos3.2 or Cy3B was accomplished using a LF405/488/561/647 quadband filter cube (TRF89902, Chroma, Bellows Falls, VT). Emission was split into two channels using a DV2 emission splitting system (Photometrics) with a T640lpxr dichroic mirror to separate emission: ET605/52m to filter near-red emission, and ET700/75m to filter far-red emission (Chroma).

Live cell imaging was accomplished at room temperature in a buffer shown previously to be compatible with BCR mediated signaling responses (30 mM Tris, 9 mg/ml glucose, 100 mM NaCl, 5 mM KCl, 1 mM KCl, 1 mM MgCl2, 1.8 mM CaCl2, 10 mM glutathione, 8 μg/ml catalase, 100 μg/ml glucose oxidase, pH 8.5 (Stone et al., 2017). Laser intensities were adjusted such that single fluorophores could be distinguished in individual frames, and were generally between 5-20 kW/cm2 for 561 nm and 647 nm lasers, and 100-200 W/cm2 for the 405 nm laser. Integration times were maintained at 20ms. Generally, 5000 frames (~2min) were acquired in unstimulated cells, then an additional 20,000 frames (8-10min) were acquired after the addition of 1 μg/ml streptavidin, which crosslinks the SiR-biotin-Fab bound to BCR and initiates a signaling response. Throughout imaging, the laser intensity is modulated to maintain a suitable density of single molecules.

For measurements involving treatment with PP2, 5 μM PP2 (Sigma: 529573) from a 10mM stock solution in DMSO was added to culture medium during the SiR-biotin-Fab labeling step prior to imaging. 5 μM PP2 was also added to all imaging buffers so that subsequent imaging and stimulation steps took place in the presence of PP2. For measurements involving treatment with alcohols, 15 μM 1-hexadecanol (Sigma: 258741) from a 6 mM DMSO stock solution or 300 μM 1-octanol (Sigma: 95446) from a 120 mM DMSO stock solution was added directly to the imaging buffer in Mattek dishes immediately before samples were transferred to the microscope for imaging. These treatments lead to a ±8C shift in miscibility transition temperatures for RBL-2H3 derived GPMVs (Gray et al., 2013; Machta et al., 2016). Final DMSO concentration was .25% for both alcohol treatments. Alcohols in DMSO were also added to all imaging buffers so that subsequent imaging and stimulation steps took place in the presence of alcohols.

Imaging of chemically fixed cells was accomplished with the same illumination and data acquisition settings in a slightly modified buffer (100 mM Tris, 9 mg/ml glucose, 25 mM NaCl, 10 mM glutathione, 8 μg/ml catalase, 100 μg/ml glucose oxidase, pH 8.5. Individual fields of view were typically imaged over 20,000 frames, and multiple fields of view were imaged in each prepared dish of cells.

### Image Processing and probe Localization

Single molecules were localized in raw live cell movies with the ImageJ plugin ThunderSTORM (Ovesný et al., 2014), using weighted least-squares fitting of an integrated Gaussian PSF with multi-emitter fitting analysis enabled to detect up to 2 single molecules within a diffraction-limited area. Localization data were then exported to our in-house MATLAB software for culling and successive post-processing steps (Veatch et al., 2012). Localizations in the near-red emission channel were registered with the far-red emission channel using a registration technique published previously (Churchman et al., 2005) and previously used by our group (Stone and Veatch, 2014, 2015). Cell regions were masked by hand for further analysis to exclude regions away from the cell as well as regions of extensive topography.

### Cross-correlation analysis

Steady-state cross-correlations from live cells were calculated as described previously (Stone and Veatch, 2015), with minor modifications. In brief, cross-correlation functions, c(r,τ), were computed from localizations in each channel that occurred in the same frame or in frames separated by a time interval τ. Long-range gradients in labeling density arising from diffusing molecules entering the TIR field along with the complicated boundary condition generated by the mask are accounted for through normalization as follows. For each channel, images reconstructed from data acquired over the entire steady-state time window are convoluted with a Gaussian kernel with σ = 1 μm and masked using the same binary mask applied to localization data. The cross-correlation of the blurred, masked images from each channel is calculated using FFTs and radially averaged as described in (Stone and Veatch, 2015; Veatch et al., 2012) and is used as a normalization factor for c(r,τ). This treatment filters structure larger than 1 μm in size from the steady-state cross-correlation function.

Bleed-through effects are evident at short τ, and this contribution is removed by fitting c(r,τ) to a sum of two Gaussian functions for each value of r:

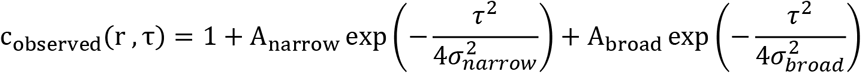

The standard deviation of the narrow Gaussian function, 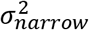, is constrained by how long it takes an anchor to diffuse across a diffraction limited distance (taken to be 250nm). This assumes that BCR is less mobile than anchors, which is consistent with experimental observations especially after BCR clustering. The broader Gaussian is interpreted to capture the more slowly varying envelope of the actual c(r, τ), most likely due to slow motions of BCR clusters. Bleed-through is accounted for by excluding the amplitude of the narrow Gaussian function to yield the values reported in the main text:

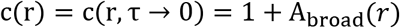

Several examples of this fitting procedure are shown in Supplementary Figure S1A. This bleed-through correction is not applied for the stimulation-time dependent curves of Fig 1D. For these measurements, the reported values are simply an average over 2s in τ:

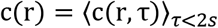

This improves statistics while reducing the impact of bleed-through in data with reduced signal to noise and produces results that are very similar to the bleed-through calculation described above, as shown in Supplementary Figure S1B.

Since we consistently observe subtle enrichment of anchors with respect to BCR in cells prior to BCR crosslinking (Supplementary Figure S2), which we attribute to long-lived membrane topography, we report on the enhanced enrichment of probes upon BCR crosslinking in order to isolate the impact of BCR clustering. This is done by subtracting c(r,τ *→* 0) obtained from images acquired prior to BCR crosslinking (t<0) from those acquired between 2-10 min after BCR crosslinking (t=2-10).

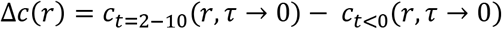

Δc is tabulated for each cell, and the variance reported in Δc represents the standard error of the mean averaged over cells. For one anchor, trLAT-2P, images were not acquired prior to BCR cross-linking for some cells included in the analysis. In this case, we subtracted the average *c*_*t*<0_(*r, τ* → 0) obtained from cells where this data was present from *c*_*t*=2-10_(*r,τ* → 0) for individual cells.

In Supplementary Figure S8, we estimate the impact of finite resolution and finite BCR cluster size on the amplitude of correlations using deconvolution, to better estimate the enrichment and depletion of anchors with respect to the centers of BCR clusters. Both resolution and the finite size of BCR clusters blur the measured Δc(r), reducing the dynamic range. Deconvolution was accomplished by first spreading Δc(r) over angles to get 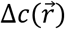 and a blurring function was estimated as a 2D Gaussian function with standard deviation of 50nm, representing the size of BCR clusters (35nm) and the approximate localization precision of mEos3.2 (35nm) added in quadrature. The 2D deconvolution was accomplished using the deconvlucy function in Matlab, using the blurring function as the point spread function, then 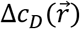 was averaged over angles to produce Δ*c_D_*(*r*). This algorithm assumes and estimates Poisson noise in the measured 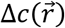. The same blurring function was used for all cells even though there is some cell to cell variation in localization precision and average BCR cluster size (Supplementary Figure S4) to simplify the analysis. A more careful correction would involve a rigorous estimate of the type and magnitude of noise from each cell and a blurring function specific to each measurement, which was not done here. The average Δ*c_D_*(*r*) curves of Figure S8 were tabulated by averaging over Δ*c_D_*(*r*) obtained from individual cells. We note that while the dynamic range of these curves increases in magnitude, so does the standard error of the mean between cells at short separation distances.

### Determination of Lo enrichment in GPMVs

RBL-2H3 or HEK cells were transiently transfected using nucleofection (Amaxa) according to protocols provided with the reagents. 4–6 h after transfection, cells were washed with PBS and then incubated with serum-free medium overnight. To synchronize the cells, 1 h before preparation of GPMVs, the cells were given full-serum medium.

Cell membranes were stained with 5 μg/ml of FAST-DiO, FAST-DiI, or DiD, respectively, green, red or far-red fluorescent lipid dyes that strongly partition to disordered phases (Levental and Levental, 2015). Following staining, GPMVs were isolated from transfected cells as described previously (Sezgin et al., 2012). GPMV formation was induced by 2 mM N-ethylmaleimide (NEM) in hypotonic buffer containing 100 mM NaCl, 10 mM HEPES, and 2 mM CaCl_2_, pH 7.4. To quantify protein partitioning, GPMVs were observed on an inverted epifluorescence microscope (Nikon) at 4°C after treatment with 200 μM DCA to stabilize phase separation; this treatment has been previously demonstrated not to affect Lo phase affinity of various proteins (Zhou et al., 2013). The relative enrichment of probes in the Lo phase is defined as (I_o_-I_d_)/(I_o_+I_d_), where I_o_ and I_d_ are baseline corrected fluorescent intensity of the construct in the liquid-ordered and liquid-disordered phases respectively for >10 vesicles/trial, with at least 3 independent experiments for each anchor. This quantity is simply related to the partition coefficient K_p_ = I_o_/I_d_ as Lo enrichment = (K_p_-1)/(K_p_+1), but is more appropriate for comparison with cell results because it is normalized to the total concentration rather than the concentration in the Lo phase, as is the case with cross-correlation functions. The majority of measurements were carried out in RBL-2H3 cells, but in some cases Hek cells were also used due to the enhanced surface expression of some probes in this cell type. When used, partitioning measurements in Hek cells produced partitioning results that were consistent with those obtained in RBL-2H3 cells but with improved signal to noise.

### Fitting to linear models and associated statistics

Cellular results were plotted vs. GPMV probe partitioning data and fit to a linear model to assess the magnitude and significance of correlations. This was done using the function lmfit() in Matlab, and in all cases values were weighted by the inverse variance of the cellular measurement. When ratios were considered, values of the stated quantity was tabulated on a cell-by-cell basis and the variance was calculated from the standard error of the mean of the ratio across cells.

The function lmfit() returns a p value against the null hypothesis that the data is described by a constant model (with no linear slope). An ANOVA analysis is then conducted using the anova() function in Matlab. The f-statistic reported is the ‘F-statistic vs. constant model’ output of this function.

In several instances, significance is reported when comparing two empirical observations. These are conducted using the ttest2 function in Matlab.

### Quantification of anchor mobility in cells

The motion of membrane anchors was quantified from localized single molecule positions using correlation methods as described previously (Stone and Veatch, 2015). Briefly, auto-correlation functions g(r, τ) were tabulated from localized positions as a function of radius r and time interval τ for each cell imaged and normalized following the same protocol as for cross-correlations described above. Correlation functions were tabulated for the same time-windows as for cross-correlation functions, either prior to BCR crosslinking (t<0) or between 2 and 10 min after activation (t=2-10). At each τ, the normalized g(r, τ) was fit to a superposition of two Gaussian functions to extract two distinct mean squared displacements according to the relation:

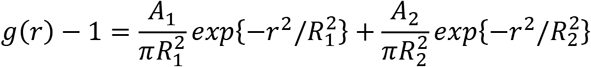

The mean squared displacement (MSD) for each component is related to the fit parameters R_1_ and R_2_ which are determined at each *τ*:

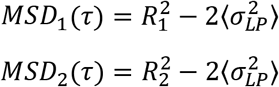

Where 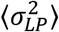 is the squared localization precision extracted from the single molecule fits and averaged over the complete dataset. The fraction of the population within the first component is *α* = *A*_1_/(*A*_1_/ + *A*_2_).

Several examples of g(r, τ) and fits are shown in Figure 5 and Sup Fig S10 and the average values of fit parameters for all probes are displayed in Sup Fig S11. We refer to the shorter MSD as the immobile component since it does not increase rapidly with τ. The longer MSD increases roughly linearly with τ, consistent with diffusive motion. We have chosen to fit to only two states for simplicity and it is likely that more mobile states are present (Karslake et al., 2020), in which case our single mobile MSD can be interpreted as a weighted average of these states. The fraction of the population within the slow component is determined at each τ and is largely independent of τ, with some exceptions (Sup Fig S10).

The MSD is related to the diffusion coefficient through the relation *MSD*(*τ*) = 4*D*(*τ*)*τ*. Since fluorophores are illuminated continuously through a finite integration time, images contain information regarding probe locations at different times throughout the integration window. As a result, the effective τ differs from the frame times of the measurement (τ_o_) (Berglund, 2010). This is accounted for as follows:

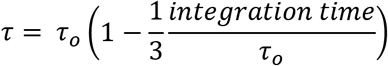

The integration time was held constant at 20ms for measurements. The frame rate (the smallest value of *τ_o_*) is slightly larger than the integration time and varies across measurements. This correction is largest for short *τ_o_* when the integration time is almost equivalent to *τ_o_*.

Motions of BCR proximal anchors were isolated from the entire population of anchors by cross-correlating anchor localizations within 100nm of a localized BCR with all anchor localizations. Since BCR clusters are largely immobile after BCR clustering, anchor localizations were included if they passed within 100nm of a BCR localization localized up to 2s previously, which implements the assumption that the BCR cluster remains fixed in space even if a fluorophore labeling BCR is not observed. Cross-correlations of BCR proximal anchors and total anchors were normalized using the same geometrical correction factor used for the cross-correlation of BCR and anchors, since the BCR proximal anchor distribution closely follows that of BCR alone. Spatio-temporal cross-correlation functions were fit to extract mobility parameters following the procedures described above for auto-correlations

### Flow cytometry

Activation of BCR through receptor cross-linking in the presence of alcohol treatments or co-clustering of BCR with minimal co-receptors (Fig 6C,D,H,I) was assessed through immunolabeling of phosphorylated BCR and measurement via flow cytometry. 500,000 CH27 cells per test condition were labeled in suspension at a concentration of 500,000 cells per ml. For alcohol treatment experiments, BCR was labeled with biotinylated goat anti-mouse IgM(μ) Fab fragments at 5 μg/ml. For co-receptor experiments, trCD4cr or trCD28cr-transduced cells were used and both BCR and co-receptors were labeled with biotinylated goat anti-mouse IgM(μ) Fab fragments at 1 μg/ml and biotinylated mouse anti-STII antibody at 5 μg/ml, respectively, for 10 min at room temperature. Unlabeled and unstimulated controls were included for all treatments. Cells were pelleted and suspended twice in PBS containing 1% BSA, then stimulated with Alexa Fluor 568-conjugated streptavidin at a final concentration of 1 μg/ml for 6 min at room temperature. For alcohol-treated samples, 15 μM 1-hexadecanol (Sigma: 258741) from a 6 mM DMSO stock solution or 300 μM 1-octanol (Sigma: 95446) from a 120mM DMSO stock solution was added directly to cells in suspension while vortexing, immediately before addition of streptavidin. Cell samples were fixed by addition of 4% paraformaldehyde to a final concentration of 1.5% and incubated for 20 min at room temperature. Cells were pelleted and rinsed with PBS-BSA to remove paraformaldehyde, then pelleted, placed on ice, and resuspended with 500 μl ice cold methanol. Cells were allowed to permeabilize on ice for 15 min, then pelleted and rinsed twice with cold PBS-BSA. Samples were resuspended in 100 μl PBS-BSA with 5 μg/ml rabbit monoclonal antibody against the CD79A subunit of BCR phosphorylated at Tyr182, and incubated for 30 min at room temperature. Cells were pelleted and resuspended twice in cold PBS-BSA to remove primary antibody, then resuspended in 200 μl PBS-BSA with 5 μg/ml Alexa Fluor 647 goat anti-rabbit IgG secondary antibody. Cells were labeled at room temperature for 60 min, then pelleted and rinsed with cold PBS-BSA. Cells were kept on ice until measurement via flow cytometry.

Flow cytometry was performed on an Attune NxT flow cytometer (ThermoFisher) with fluorescence measurements of eGFP, Alexa Fluor 568, and Alexa Fluor 647 using 488, 561, and 647nm laser lines for co-receptor experiments or Alexa Fluor 568 and Alexa Fluor 647 using 561 and 647nm laser lines for alcohol treatment experiments. Fluorescence bleed-through was corrected using compensation procedures. Data was exported as .fcs files and was read and analyzed using the fca_readfcs.m function (Balkay, 2020) and custom analysis scripts in Matlab. Single intact cells were gated based on forward- and back-scatter measurements. For trCD4cr and trCD28cr expressing cells, cell populations were gated based on a narrow range of eGFP fluorescence so that the pBCR signal was compared for cells expressing equivalent levels of trCD4cr vs. trCD28cr.

### Generation of the membrane for Figure 7

The membrane shown in Fig 7 was generated from a 2D matrix of pixel values (p), half normally distributed around the value −1 and half normally distributed around the value +1. Pixels were allowed to redistribute across the matrix through non-local exchanges and with periodic boundary conditions using a Monte Carlo algorithm that satisfied detailed balance, which tabulates the energy at each position i as 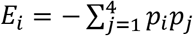 where *p_j_* represents the pixel value of the 4 nearest neighbors. Temperature was chosen such that the size of structure in the simulation was small compared to the size of the matrix.

After equilibration, a weak field was introduced to bias the location of pixels within certain regions of the matrix. This field provided the additional energy *Ef_i_* = *p_i_F_i_*. Plots showing the distribution of pixel values and the applied field matrix are found in Supplementary Figure S16. The matrix was square (400 by 400 pixels) and the field was applied in a region of the simulation. The remaining values of the field were set to zero. The average image was generated by averaging 50 different simulation snapshots with the same applied field.

## MULTIMEDIA FILES

**Supplementary Movie 1: Single molecule motions of BCR and PM before BCR crosslinking.** Time-lapse video showing reconstructed single molecule positions evolving in time for BCR (magenta) and PM (green) for the same PM expressing cell shown in Fig 1. A-C. These localizations were acquired <1min prior to the addition of streptavidin. Localizations are drawn on a reconstructed image of all PM localizations for this cell. The movie is displayed with a frame rate that is 5x slower than real time. Scale bar is 5 μm.

**Supplementary Movie 2: Single molecule motions of BCR and PM after BCR crosslinking.** Time-lapse video showing reconstructed single molecule positions evolving in time for BCR (magenta) and PM (green) for the same PM expressing cell shown in Fig 1. A-C. These localizations were acquired ~2min after addition of streptavidin. Localizations are drawn on a reconstructed image of all PM localizations for this cell. The movie is displayed with a frame rate that is 5x slower than real time. Scale bar is 5 μm.

**Supplementary Movie 3: Single molecule motions of BCR and M before BCR crosslinking.** Time-lapse video showing reconstructed single molecule positions evolving in time for BCR (magenta) and M (green) for the same M expressing cell shown in Fig 1. A-C. These localizations were acquired <1min prior to the addition of streptavidin. Localizations are drawn on a reconstructed image of all M localizations for this cell. The movie is displayed with a frame rate that is 5x slower than real time. Scale bar is 5 μm.

**Supplementary Movie 4: Single molecule motions of BCR and M after BCR crosslinking.** Time-lapse video showing reconstructed single molecule positions evolving in time for BCR (magenta) and M (green) for the same M expressing cell shown in Fig 1. A-C. These localizations were acquired ~2min after addition of streptavidin. Localizations are drawn on a reconstructed image of all M localizations for this cell. The movie is displayed with a frame rate that is 5x slower than real time. Scale bar is 5 μm.

**Supplementary Movie 5: Time lapse of BCR and PM organization before and after BCR cross-linking.** Time lapse movie composed of reconstructed images of BCR (magenta) and PM (green) from the same representative cell shown in Fig 1. A-C, shown here at higher time resolution to demonstrate how BCR and PM organization evolve over time. Each sequential image is reconstructed from a window of 750 frames (~20sec) incremented by 250 frames for each image. The entire time lapse spans the length of the live cell experiment and the real time of data acquisition is shown in the bottom-left, where 0 min corresponds to streptavidin addition. Before streptavidin addition, both BCR and PM are relatively uniformly distributed across the plasma membrane. After streptavidin addition, BCR rapidly forms tight clusters. BCR clusters are relatively immobile compared to PM, but do exhibit dynamics that are clearly evident in the time lapse. Scale bar is 5 μm.

**Supplementary Movie 6: Time lapse of BCR and M organization before and after BCR cross-linking.** Time lapse movie composed of reconstructed images of BCR (magenta) and M (green)from the same representative cell shown in Fig 1. AC. Parameters for reconstruction of individual frames of the time lapse are the same as in Supplementary Movie 3, and the time lapse shows similar organization of BCR and M before and after streptavidin addition. Time before and after streptavidin addition in shown in the bottom left and the scale bar is 5 μm.

## Notes

https://sites.lsa.umich.edu/veatch-lab/science-images/

### Competing Interest Statement

The authors have declared no competing interest.

### Summary of Updates

Results section is updated to include a new figure (Fig 6) showing the effect of perturbations on probe partitioning and BCR tyrosine phosphorylation. The manuscript has been revised for clarity throughout.

## REFERENCES

A. Camley, B., and H. Brown, F.L. (2013). Diffusion of complex objects embedded in free and supported lipid bilayer membranes: role of shape anisotropy and leaflet structure. Soft Matter 9, 4767–4779.

Arumugam, S., Petrov, E.P., and Schwille, P. (2015). Cytoskeletal Pinning Controls Phase Separation in Multicomponent Lipid Membranes. Biophys. J. 108, 1104–1113.

Bag, N., Wagenknecht-Wiesner, A., Lee, A., Shi, S.M., Holowka, D.A., and Baird, B.A. (2021). Lipid-based and protein-based interactions synergize transmembrane signaling stimulated by antigen clustering of IgE receptors. Proc. Natl. Acad. Sci. U. S. A. 118, e2026583118.

Balkay, L. (2020). fca_readfcs (https://www.mathworks.com/matlabcentral/fileexchange/9608-fca_readfcs), MATLAB Central File Exchange.

Baumgart, T., Hammond, A.T., Sengupta, P., Hess, S.T., Holowka, D.A., Baird, B.A., and Webb, W.W. (2007). Large-scale fluid/fluid phase separation of proteins and lipids in giant plasma membrane vesicles. Proc. Natl. Acad. Sci. U. S. A. 104, 3165–3170.

Berglund, A.J. (2010). Statistics of camera-based single-particle tracking. Phys. Rev. E 82, 011917.

Bloom, M., and Thewalt, J.L. (1995). Time and distance scales of membrane domain organization. Mol. Membr. Biol. 12, 9–13.

Brown, D.A., and Rose, J.K. (1992). Sorting of GPI-anchored proteins to glycolipid-enriched membrane subdomains during transport to the apical cell surface. Cell 68, 533–544.

Burns, M., Wisser, K., Wu, J., Levental, I., and Veatch, S.L. (2017). Miscibility Transition Temperature Scales with Growth Temperature in a Zebrafish Cell Line. Biophys. J. 113, 1212–1222.

Cammarota, E., Soriani, C., Taub, R., Morgan, F., Sakai, J., Veatch, S.L., Bryant, C.E., and Cicuta, P. (2020). Criticality of plasma membrane lipids reflects activation state of macrophage cells. J. R. Soc. Interface 17, 20190803.

Cherukuri, A., Cheng, P.C., Sohn, H.W., and Pierce, S.K. (2001). The CD19/CD21 Complex Functions to Prolong B Cell Antigen Receptor Signaling from Lipid Rafts. Immunity 14, 169–179.

Cherukuri, A., Carter, R.H., Brooks, S., Bornmann, W., Finn, R., Dowd, C.S., and Pierce, S.K. (2004). B cell signaling is regulated by induced palmitoylation of CD81. J. Biol. Chem. 279, 31973–31982.

Chung, J.K., Huang, W.Y.C., Carbone, C.B., Nocka, L.M., Parikh, A.N., Vale, R.D., and Groves, J.T. (2021). Coupled membrane lipid miscibility and phosphotyrosine-driven protein condensation phase transitions. Biophys. J. 120, 1257–1265.

Churchman, L.S., Ökten, Z., Rock, R.S., Dawson, J.F., and Spudich, J.A. (2005). Single molecule high-resolution colocalization of Cy3 and Cy5 attached to macromolecules measures intramolecular distances through time. Proc. Natl. Acad. Sci. U. S. A. 102, 1419–1423.

Dart, C. (2010). SYMPOSIUM REVIEW: Lipid microdomains and the regulation of ion channel function. J. Physiol. 588, 3169–3178.

Davis, S.J., and van der Merwe, P.A. (2006). The kinetic-segregation model: TCR triggering and beyond. Nat. Immunol. 7, 803–809.

Delos Santos, R.C., Garay, C., and Antonescu, C.N. (2015). Charming neighborhoods on the cell surface: Plasma membrane microdomains regulate receptor tyrosine kinase signaling. Cell. Signal. 27, 1963–1976.

Diaz-Rohrer, B.B., Levental, K.R., Simons, K., and Levental, I. (2014). Membrane raft association is a determinant of plasma membrane localization. Proc. Natl. Acad. Sci. U. S. A. 111, 8500–8505.

Eggeling, C., Ringemann, C., Medda, R., Schwarzmann, G., Sandhoff, K., Polyakova, S., Belov, V.N., Hein, B., von Middendorff, C., Schönle, A., et al. (2009). Direct observation of the nanoscale dynamics of membrane lipids in a living cell. Nature 457, 1159–1162.

Elliott, J.I., Sardini, A., Cooper, J.C., Alexander, D.R., Davanture, S., Chimini, G., and Higgins, C.F. (2006). Phosphatidylserine exposure in B lymphocytes: a role for lipid packing. Blood 108, 1611–1617.

Ernst, R., Ejsing, C.S., and Antonny, B. (2016). Homeoviscous Adaptation and the Regulation of Membrane Lipids. J. Mol. Biol. 428, 4776–4791.

Fujiwara, T., Ritchie, K., Murakoshi, H., Jacobson, K., and Kusumi, A. (2002). Phospholipids undergo hop diffusion in compartmentalized cell membrane. J. Cell Biol. 157, 1071–1082.

Ganti, R.S., Lo, W.-L., McAffee, D.B., Groves, J.T., Weiss, A., and Chakraborty, A.K. (2020). How the T cell signaling network processes information to discriminate between self and agonist ligands. Proc. Natl. Acad. Sci. 117, 26020–26030.

Gasparrini, F., Feest, C., Bruckbauer, A., Mattila, P.K., Müller, J., Nitschke, L., Bray, D., and Batista, F.D. (2016). Nanoscale organization and dynamics of the siglec CD22 cooperate with the cytoskeleton in restraining BCR signalling. EMBO J. 35, 258–280.

Gauld, S.B., Porto, J.M.D., and Cambier, J.C. (2002). B Cell Antigen Receptor Signaling: Roles in Cell Development and Disease. Science 296, 1641–1642.

Ge, M., Gidwani, A., Brown, H.A., Holowka, D., Baird, B., and Freed, J.H. (2003). Ordered and Disordered Phases Coexist in Plasma Membrane Vesicles of RBL-2H3 Mast Cells. An ESR Study. Biophys. J. 85, 1278–1288.

Gold, M.R., and Reth, M.G. (2019). Antigen Receptor Function in the Context of the Nanoscale Organization of the B Cell Membrane. Annu. Rev. Immunol. 37, 97–123.

Gray, E., Karslake, J., Machta, B.B., and Veatch, S.L. (2013). Liquid General Anesthetics Lower Critical Temperatures in Plasma Membrane Vesicles. Biophys. J. 105, 2751–2759.

Guigas, G., and Weiss, M. (2016). Effects of protein crowding on membrane systems. Biochim. Biophys. Acta BBA - Biomembr. 1858, 2441–2450.

Gupta, N., and DeFranco, A.L. (2007). Lipid rafts and B cell signaling. Semin. Cell Dev. Biol. 18, 616–626.

Hanzal-Bayer, M.F., and Hancock, J.F. (2007). Lipid rafts and membrane traffic. FEBS Lett. 581, 2098–2104.

Haughton, G., Arnold, L.W., Bishop, G.A., and Mercolino, T.J. (1986). The CH Series of Murine B Cell Lymphomas: Neoplastic Analogues of Ly-1+ Normal B Cells. Immunol. Rev. 93, 35–52.

Heberle, F.A., Wu, J., Goh, S.L., Petruzielo, R.S., and Feigenson, G.W. (2010). Comparison of Three Ternary Lipid Bilayer Mixtures: FRET and ESR Reveal Nanodomains. Biophys. J. 99, 3309–3318.

Heberle, F.A., Doktorova, M., Scott, H.L., Skinkle, A.D., Waxham, M.N., and Levental, I. (2020). Direct label-free imaging of nanodomains in biomimetic and biological membranes by cryogenic electron microscopy. Proc. Natl. Acad. Sci. 117, 19943–19952.

Heinrich, M., Tian, A., Esposito, C., and Baumgart, T. (2010). Dynamic sorting of lipids and proteins in membrane tubes with a moving phase boundary. Proc. Natl. Acad. Sci. 107, 7208–7213.

Holowka, D., and Baird, B. (2016). Roles for lipid heterogeneity in immunoreceptor signaling. Biochim. Biophys. Acta 1861, 830–836.

Honigmann, A., Sadeghi, S., Keller, J., Hell, S.W., Eggeling, C., and Vink, R. (2014). A lipid bound actin meshwork organizes liquid phase separation in model membranes. ELife 3.

Igarashi, M., Honda, A., Kawasaki, A., and Nozumi, M. (2020). Neuronal Signaling Involved in Neuronal Polarization and Growth: Lipid Rafts and Phosphorylation. Front. Mol. Neurosci. 13.

Ingólfsson, H.I., Melo, M.N., van Eerden, F.J., Arnarez, C., Lopez, C.A., Wassenaar, T.A., Periole, X., de Vries, A.H., Tieleman, D.P., and Marrink, S.J. (2014). Lipid Organization of the Plasma Membrane. J. Am. Chem. Soc. 136, 14554–14559.

Kabbani, A.M., Raghunathan, K., Lencer, W.I., Kenworthy, A.K., and Kelly, C.V. (2020). Structured clustering of the glycosphingolipid GM1 is required for membrane curvature induced by cholera toxin. Proc. Natl. Acad. Sci. 117, 14978–14986.

Kalland, M.E., Solheim, S.A., Skånland, S.S., Taskén, K., and Berge, T. (2012). Modulation of proximal signaling in normal and transformed B cells by transmembrane adapter Cbp/PAG. Exp. Cell Res. 318, 1611–1619.

Karnell, F.G., Brezski, R.J., King, L.B., Silverman, M.A., and Monroe, J.G. (2005). Membrane cholesterol content accounts for developmental differences in surface B cell receptor compartmentalization and signaling. J. Biol. Chem. 280, 25621–25628.

Karslake, J.D., Donarski, E.D., Shelby, S.A., Demey, L.M., DiRita, V.J., Veatch, S.L., and Biteen, J.S. (2020). SMAUG: Analyzing single-molecule tracks with nonparametric Bayesian statistics. Methods.

Kenworthy, A.K. (2008). Have we become overly reliant on lipid rafts? Talking Point on the involvement of lipid rafts in T-cell activation. EMBO Rep. 9, 531–535.

Kenworthy, A.K., Nichols, B.J., Remmert, C.L., Hendrix, G.M., Kumar, M., Zimmerberg, J., and Lippincott-Schwartz, J. (2004). Dynamics of putative raft-associated proteins at the cell surface. J. Cell Biol. 165, 735–746.

Kinoshita, M., Suzuki, K.G.N., Matsumori, N., Takada, M., Ano, H., Morigaki, K., Abe, M., Makino, A., Kobayashi, T., Hirosawa, K.M., et al. (2017). Raft-based sphingomyelin interactions revealed by new fluorescent sphingomyelin analogs. J. Cell Biol. 216, 1183–1204.

Knight, J.D., Lerner, M.G., Marcano-Velázquez, J.G., Pastor, R.W., and Falke, J.J. (2010). Single Molecule Diffusion of Membrane-Bound Proteins: Window into Lipid Contacts and Bilayer Dynamics. Biophys. J. 99, 2879–2887.

Koyama-Honda, I., Fujiwara, T.K., Kasai, R.S., Suzuki, K.G.N., Kajikawa, E., Tsuboi, H., Tsunoyama, T.A., and Kusumi, A. (2020). High-speed single-molecule imaging reveals signal transduction by induced transbilayer raft phases. J. Cell Biol. 219.

Kraft, M.L. (2013). Plasma membrane organization and function: moving past lipid rafts. Mol. Biol. Cell 24, 2765–2768.

Lee, I.-H., Imanaka, M.Y., Modahl, E.H., and Torres-Ocampo, A.P. (2019). Lipid Raft Phase Modulation by Membrane-Anchored Proteins with Inherent Phase Separation Properties. ACS Omega 4, 6551–6559.

Levental, K.R., and Levental, I. (2015). Isolation of Giant Plasma Membrane Vesicles for Evaluation of Plasma Membrane Structure and Protein Partitioning. In Methods in Membrane Lipids, D.M. Owen, ed. (New York, NY: Springer), pp. 65–77.

Levental, I., Lingwood, D., Grzybek, M., Coskun, U., and Simons, K. (2010). Palmitoylation regulates raft affinity for the majority of integral raft proteins. Proc. Natl. Acad. Sci. U. S. A. 107, 22050–22054.

Levental, I., Levental, K.R., and Heberle, F.A. (2020a). Lipid Rafts: Controversies Resolved, Mysteries Remain. Trends Cell Biol. 30, 341–353.

Levental, K.R., Surma, M.A., Skinkle, A.D., Lorent, J.H., Zhou, Y., Klose, C., Chang, J.T., Hancock, J.F., and Levental, I. (2017). ω-3 polyunsaturated fatty acids direct differentiation of the membrane phenotype in mesenchymal stem cells to potentiate osteogenesis. Sci. Adv. 3, eaao1193.

Levental, K.R., Malmberg, E., Symons, J.L., Fan, Y.-Y., Chapkin, R.S., Ernst, R., and Levental, I. (2020b). Lipidomic and biophysical homeostasis of mammalian membranes counteracts dietary lipid perturbations to maintain cellular fitness. Nat. Commun. 11, 1339.

Li, G., Wang, Q., Kakuda, S., and London, E. (2020). Nanodomains can persist at physiologic temperature in plasma membrane vesicles and be modulated by altering cell lipids[S]. J. Lipid Res. 61, 758–766.

Lingwood, D., and Simons, K. (2010). Lipid rafts as a membrane-organizing principle. Science 327, 46–50.

Liu, A.P., and Fletcher, D.A. (2006). Actin Polymerization Serves as a Membrane Domain Switch in Model Lipid Bilayers. Biophys. J. 91, 4064–4070.

Liu, L., Sommermeyer, D., Cabanov, A., Kosasih, P., Hill, T., and Riddell, S.R. (2016). Inclusion of Strep-tag II in design of antigen receptors for T-cell immunotherapy. Nat. Biotechnol. 34, 430–434.

Lorent, J.H., Diaz-Rohrer, B., Lin, X., Spring, K., Gorfe, A.A., Levental, K.R., and Levental, I. (2017). Structural determinants and functional consequences of protein affinity for membrane rafts. Nat. Commun. 8, 1219.

Lyon, A.S., Peeples, W.B., and Rosen, M.K. (2021). A framework for understanding the functions of biomolecular condensates across scales. Nat. Rev. Mol. Cell Biol. 22, 215–235.

Machta, B.B., Papanikolaou, S., Sethna, J.P., and Veatch, S.L. (2011). Minimal Model of Plasma Membrane Heterogeneity Requires Coupling Cortical Actin to Criticality. Biophys. J. 100, 1668–1677.

Machta, B.B., Veatch, S.L., and Sethna, J.P. (2012). Critical Casimir Forces in Cellular Membranes. Phys. Rev. Lett. 109, 138101.

Machta, B.B., Gray, E., Nouri, M., McCarthy, N.L.C., Gray, E.M., Miller, A.L., Brooks, N.J., and Veatch, S.L. (2016). Conditions that Stabilize Membrane Domains Also Antagonize n-Alcohol Anesthesia. Biophys. J. 111, 537–545.

Manley, S., Horton, M.R., Lecszynski, S., and Gast, A.P. (2008). Sorting of Streptavidin Protein Coats on Phase-Separating Model Membranes. Biophys. J. 95, 2301–2307.

Marshall, A.J., Niiro, H., Yun, T.J., and Clark, E.A. (2000). Regulation of B-cell activation and differentiation by the phosphatidylinositol 3-kinase and phospholipase Cgamma pathway. Immunol. Rev. 176, 30–46.

Mattila, P.K., Batista, F.D., and Treanor, B. (2016). Dynamics of the actin cytoskeleton mediates receptor cross talk: An emerging concept in tuning receptor signaling. J. Cell Biol. 212, 267–280.

McKeithan, T.W. (1995). Kinetic proofreading in T-cell receptor signal transduction. Proc. Natl. Acad. Sci. 92, 5042–5046.

Munro, S. (2003). Lipid Rafts: Elusive or Illusive? Cell 115, 377–388.

Oldfield, E., and Chapman, D. (1972). Dynamics of lipids in membranes: Heterogeneity and the role of cholesterol. FEBS Lett. 23, 285–297.

Ovesný, M., Křížek, P., Borkovec, J., Svindrych, Z., and Hagen, G.M. (2014). ThunderSTORM: a comprehensive ImageJ plug-in for PALM and STORM data analysis and super-resolution imaging. Bioinforma. Oxf. Engl. 30, 2389–2390.

Packard, T.A., and Cambier, J.C. (2013). B lymphocyte antigen receptor signaling: initiation, amplification, and regulation. F1000prime Rep. 5, 40.

Pathak, P., and London, E. (2015). The Effect of Membrane Lipid Composition on the Formation of Lipid Ultrananodomains. Biophys. J. 109, 1630–1638.

Pierce, S.K., and Liu, W. (2010). The tipping points in the initiation of B cell signalling: how small changes make big differences. Nat. Rev. Immunol. 10, 767–777.

Pore, D., and Gupta, N. (2015). Ezrin-Radixin-Moesin family proteins in the regulation of B cell immune response. Crit. Rev. Immunol. 35, 15–31.

Saffman, P.G., and Delbrück, M. (1975). Brownian motion in biological membranes. Proc. Natl. Acad. Sci. 72, 3111–3113.

Sanchez, S.A., Tricerri, M.A., and Gratton, E. (2012). Laurdan generalized polarization fluctuations measures membrane packing micro-heterogeneity in vivo. Proc. Natl. Acad. Sci. U. S. A. 109, 7314–7319.

Schmid, F. (2017). Physical mechanisms of micro- and nanodomain formation in multicomponent lipid membranes. Biochim. Biophys. Acta BBA - Biomembr. 1859, 509–528.

Schneider, F., Waithe, D., Clausen, M.P., Galiani, S., Koller, T., Ozhan, G., Eggeling, C., and Sezgin, E. (2017). Diffusion of lipids and GPI-anchored proteins in actin-free plasma membrane vesicles measured by STED-FCS. Mol. Biol. Cell 28, 1507–1518.

Sengupta, P., Holowka, D., and Baird, B. (2007). Fluorescence resonance energy transfer between lipid probes detects nanoscopic heterogeneity in the plasma membrane of live cells. Biophys. J. 92, 3564–3574.

Sezgin, E., Kaiser, H.-J., Baumgart, T., Schwille, P., Simons, K., and Levental, I. (2012). Elucidating membrane structure and protein behavior using giant plasma membrane vesicles. Nat. Protoc. 7, 1042–1051.

Shaw, T.R., Ghosh, S., and Veatch, S.L. (2021). Critical Phenomena in Plasma Membrane Organization and Function. Annu. Rev. Phys. Chem. 72, null.

Shelby, S.A., Veatch, S.L., Holowka, D.A., and Baird, B.A. (2016). Functional nanoscale coupling of Lyn kinase with IgE-FcεRI is restricted by the actin cytoskeleton in early antigen-stimulated signaling. Mol. Biol. Cell 27, 3645–3658.

Shin, Y., and Brangwynne, C.P. (2017). Liquid phase condensation in cell physiology and disease. Science 357.

Simons, K., and Ikonen, E. (1997). Functional rafts in cell membranes. Nature 387, 569–572.

Sohn, H.W., Tolar, P., Jin, T., and Pierce, S.K. (2006). Fluorescence resonance energy transfer in living cells reveals dynamic membrane changes in the initiation of B cell signaling. Proc. Natl. Acad. Sci. U. S. A. 103, 8143–8148.

Sorre, B., Callan-Jones, A., Manneville, J.-B., Nassoy, P., Joanny, J.-F., Prost, J., Goud, B., and Bassereau, P. (2009). Curvature-driven lipid sorting needs proximity to a demixing point and is aided by proteins. Proc. Natl. Acad. Sci. 106, 5622–5626.

Sproul, T.W., Malapati, S., Kim, J., and Pierce, S.K. (2000). Cutting Edge: B Cell Antigen Receptor Signaling Occurs Outside Lipid Rafts in Immature B Cells. J. Immunol. 165, 6020–6023.

St. Clair, J.W., Kakuda, S., and London, E. (2020). Induction of Ordered Lipid Raft Domain Formation by Loss of Lipid Asymmetry. Biophys. J. 119, 483–492.

Stone, M.B., and Veatch, S.L. (2014). Far-Red Organic Fluorophores Contain a Fluorescent Impurity. ChemPhysChem 15, 2240–2246.

Stone, M.B., and Veatch, S.L. (2015). Steady-state cross-correlations for live two-colour super-resolution localization data sets. Nat. Commun. 6.

Stone, M.B., Shelby, S.A., Núñez, M.F., Wisser, K., and Veatch, S.L. (2017). Protein sorting by lipid phase-like domains supports emergent signaling function in B lymphocyte plasma membranes. ELife 6, e19891.

Strzyz, P. (2019). Tying lipid rafts to oncogenic signalling. Nat. Rev. Mol. Cell Biol. 20, 513.

Sviridov, D., and Miller, Y.I. (2020). Biology of Lipid Rafts: Introduction to the Thematic Review Series. J. Lipid Res. 61, 598–600.

Tischer, D.K., and Weiner, O.D. (2019). Light-based tuning of ligand half-life supports kinetic proofreading model of T cell signaling. ELife 8, e42498.

Tolar, P., Hanna, J., Krueger, P.D., and Pierce, S.K. (2009). The constant region of the membrane immunoglobulin mediates B cell-receptor clustering and signaling in response to membrane antigens. Immunity 30, 44–55.

Varshney, P., Yadav, V., and Saini, N. (2016). Lipid rafts in immune signalling: current progress and future perspective. Immunology 149, 13–24.

Veatch, S.L., Cicuta, P., Sengupta, P., Honerkamp-Smith, A., Holowka, D., and Baird, B. (2008). Critical Fluctuations in Plasma Membrane Vesicles. ACS Chem. Biol. 3, 287–293.

Veatch, S.L., Machta, B.B., Shelby, S.A., Chiang, E.N., Holowka, D.A., and Baird, B.A. (2012). Correlation Functions Quantify Super-Resolution Images and Estimate Apparent Clustering Due to Over-Counting. PLoS ONE 7, e31457.

Vogel, S.K., Greiss, F., Khmelinskaia, A., and Schwille, P. (2017). Control of lipid domain organization by a biomimetic contractile actomyosin cortex. ELife 6, e24350.

Yam, P.Y., Li, S., Wu, J., Hu, J., Zaia, J.A., and Yee, J.-K. (2002). Design of HIV Vectors for Efficient Gene Delivery into Human Hematopoietic Cells. Mol. Ther. 5, 479–484.

Yousefi, O.S., Günther, M., Hörner, M., Chalupsky, J., Wess, M., Brandl, S.M., Smith, R.W., Fleck, C., Kunkel, T., Zurbriggen, M.D., et al. (2019). Optogenetic control shows that kinetic proofreading regulates the activity of the T cell receptor. ELife 8, e42475.

Zhao, J., Wu, J., and Veatch, S.L. (2013). Adhesion Stabilizes Robust Lipid Heterogeneity in Supercritical Membranes at Physiological Temperature. Biophys. J. 104, 825–834.

Zhou, Y., Maxwell, K.N., Sezgin, E., Lu, M., Liang, H., Hancock, J.F., Dial, E.J., Lichtenberger, L.M., and Levental, I. (2013). Bile Acids Modulate Signaling by Functional Perturbation of Plasma Membrane Domains. J. Biol. Chem. 288, 35660–35670.

