## Supporting table and figures for "Membrane phase separation drives organization at B cell receptor clusters"

| Anchor name | Description | Sequence | Source/citation |
|---|---|---|---|
| trLAT | Transmembrane domain from human LAT. mEos3.2 fused on the C terminus. | N'-MEEAILVPCVLGLLLLPILAMLMALCVHCHRLPGSGIH-[mEos3.2]-C' | Ilya Levental Lab [1] |
| trLATo | Transmembrane domain from mouse LAT. mEos3.2 fused on the N terminus. Also contains an N-terminal ER translocation signal. | N'-[ER]-ARDPPVAT-[mEos3.2]-FSRMEADALSPVGLGLLLLPFLVTLLAALCVRCRELPVS-C' | Jonathan Grover – Akira Ono Lab |
| trLAT-P | Identical to trLATo with the palmitoylation site at C27 mutated to A. | N'-[ER]-ARDPPVAT-[mEos3.2]-FSRMEADALSPVGLGLLLLPFLVTLLAAL**A**VRCRELPVS-C' | Jonathan Grover – Akira Ono Lab |
| trLAT-2P | Identical to trLATo with both palmitoylation sites at C27 and C30 mutated to A. | N'-[ER]-ARDPPVAT-[mEos3.2]-FSRMEADALSPVGLGLLLLPFLVTLLAAL**A**VR**A**RELPVS-C' | Jonathan Grover – Akira Ono Lab [2,3] |
| trLATm 1 (PtoL) | Identical to trLAT with prolines P8 and P17 in the LAT transmembrane domain mutated to L. | N'-MEEAILVLCVLGLLLLLILAMLMALCVHCHRLPGSGIH-[mEos3.2]-C' | Ilya Levental Lab [4] |
| trLATm 2 (allL+A) | Identical to trLAT with residues within the transmembrane domain mutated to L and A. This modification increases the TMD interfacial surface area with the membrane [4]. | N'-MEELLALLLLALAAAALLALLALLACVHCHRLPGSGIH-[mEos3.2]-C' | Ilya Levental Lab [4] |
| trCD4 (trLAT/ CD4) | Transmembrane domain from CD4, contains 2 palmitoylation sites. mEos3.2 fused on the C terminus. | N'-MEEVQPMALIVLGGVAGLLLFIGLGIFFCVHCHRLPGSGIH-[mEos3.2]-C' | Ilya Levental Lab [4,5] |
| trCD4o | Transmembrane domain from CD4, contains 2 palmitoylation sites. mEos3.2 fused on the N terminus. | N'-[mEos3.2]-FSSSSFEFMESNIKVLPTWSTPVQPMALIVLGGVAGLLLFIGLGIFFCVRCRHRRRQGSTGSR-C' | Sarah Veatch Lab (unpublished) |
| trPAG | Transmembrane domain from PAG/CSK, contains 2 palmitoylation sites. mEos3.2 fused on the C terminus. | N'-MQITLWGSLAAVAIFFVITFLIFLCSSCHRLPGSGIH-[mEos3.2]-C' | Ilya Levental Lab [4,5] |
| trCD45 | Transmembrane domain from CD45. mEos3.2 fused on the N terminus. | N'-[mEos3.2]-SRGTMMTPNESTNFNAKALIIFLVFLIIVTSIALLVVLYKIYDLRKKRADPPDLDN-C' | Sarah Veatch Lab (unpublished) |
| M | N terminus of Src with 1 myristoylation and short polybasic sequence. | N'-MGSSKSKPKDPSQRRNNNNGPVAT-[mEos3.2]-C' | William Rodgers Lab [6] |
| PM | N terminus of Lyn with 1 myristoylation and 1 palmitoylation. | N'-MGCIKSKRKDKDLELKLRILQSTVPRARDPPVAT-[mEos3.2]-C' | Barbara Baird and David Holowka Lab [7,8] |
| PPM | N terminus of Fyn with 1 myristoylation and 2 palmitoylations. | N'-MGCVQCKDKE-[mEos3.2]-C' | Akira Ono Lab [9] |
| GG | C terminus of KRas with polybasic sequence and original farnesylation replaced by geranylgeranylation. | N'-[mEos3.2]-FRSDGKKKKKKSKTKCQLL-C' | Barbara Baird and David Holowka Lab [7,8] |
| FP | C terminus of NRas with 1 farnesylationand 1 palmitoylation. | N'-[mEos3.2]-FSSSSLNSAVDGCMGLPCVVM-C' | John Hancock Lab [10,11] |
| FPP | C terminus of HRas with 1 farnesylation and 2 palmitoylations. | N'-[mEos3.2]-FSSSSLNSGCMSCKCVLS-C' | John Hancock Lab [10,11] |
| GPI | C-terminal sequence derived from CD58 encoding a GPI-attachment signal. | N'-[mEos3.2]-YGGNGSGQHQYDPRPSSGHSRHRYALIPIPLAVITTCIVLYMNVL-C' | Kai Simons Lab [12] |

**Supplementary Table 1: Sequence, design, and origin information for anchor constructs.** Descriptions, sequences, sources and references (if available) are provided for the anchor constructs used in this study. For GPMV measurements, cells expressed anchors conjugated to either GFP, YFP, or RFP, as described in the original publications. For FLM measurements, anchors are conjugated to mEos3.2. The full sequence of mEos3.2 is omitted in the table but listed below. Some anchor constructs contain ER translocation signal sequences that improve delivery to the plasma membrane. The ER translocation signal sequence is also omitted from the table but is listed below. Underlined sections of sequences represent portions taken from the native protein. Other parts of the sequence are linkers or overhangs used for cloning.

*mEos3.2 sequence [mEos3.2]:*
MSAIKPDMKIKLRMEGNVNGHHFVIDGDGTGKPFEGKQSMDLEVKEGGPLPFAFDILTTAFHYGNRVFAKYPDNIQDYFKQSFPKGYSWERSLTFEDGGICNARNDITMEGDTFYNKVRFYGTNFPANGPVMQKKTLKWEPSTEKMYVRDGVLTGDIEMALLLEGNAHYRCDFRTTYKAKEKGVKLPGAHFVDHCIEILSHDKDYNKVKLYEHAVAHSGLPDNARR

*ER translocation sequence [ER]:* MELFWSIVFTVLLSFSCRGSDWESLQSTVPR

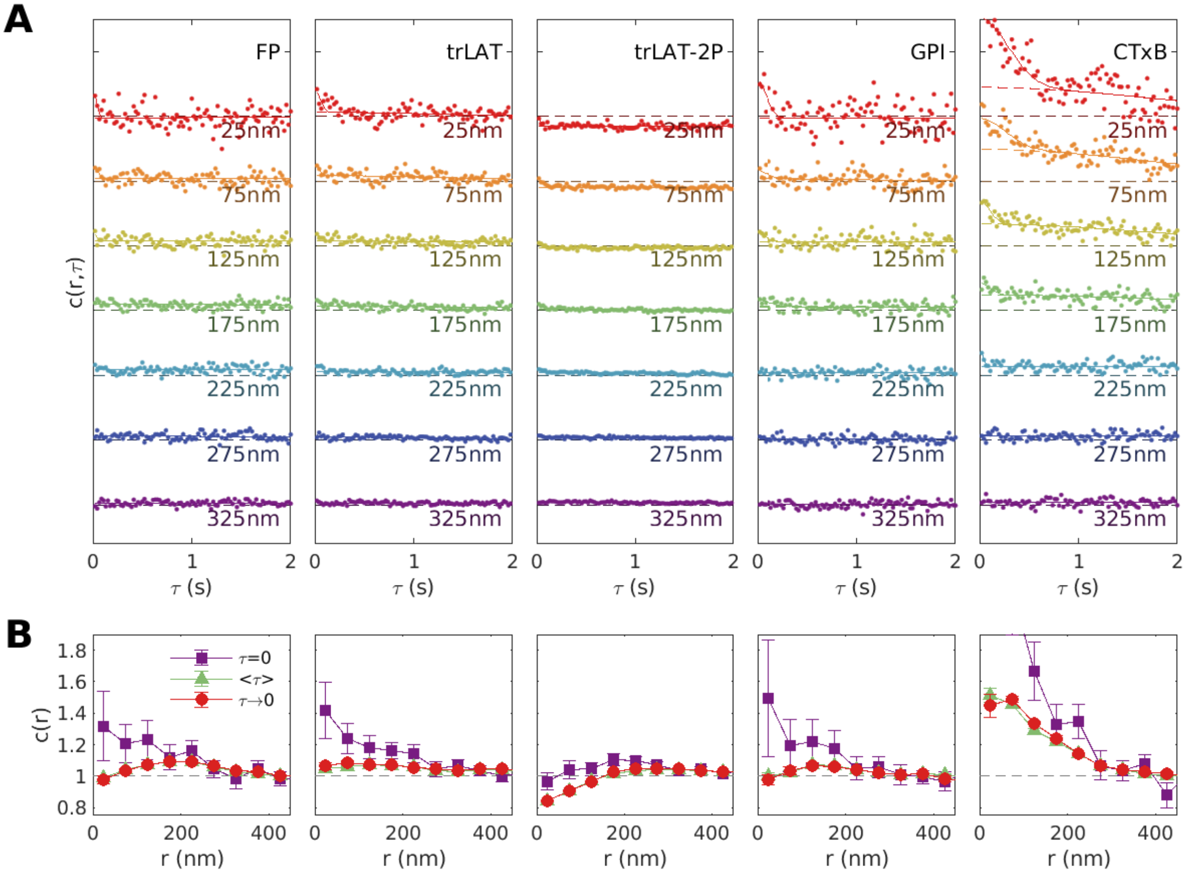


**Supplementary Figure S1: Considering finite time-intervals (τ) improves counting statistics and corrects for cross-talk between imaging channels.** (A) Raw c(r, τ) curves tabulated from single cells viewed 2-10min after BCR crosslinking. Points are fit to a superposition of two Gaussian functions (solid lines) to capture amplitude variation that occurs on distinct time-scales. We interpret long-time variation as arising from slow motions of BCR clusters. Short-time variation is due to bleed-through between color channels and its range is bounded by probe diffusion time through a diffraction limited area. This bleed-through effect is most prominent for CTxB because this is the only anchor probed that is labeled with an organic fluorophore (Cy3B), which is brighter than mEos3.2 and bleeds more into the SiR (far red) emission channel. CTxB is also the slowest anchor probed, so bleed-through extends to larger τ. Bleed-through is corrected by extrapolating the long-range Gaussian fit to τ=0 (dashed lines), and we refer to this as c(r, τ→0). Since these curves are determined by many points (~100), the extrapolated c(r, τ→0) is determined with smaller error bounds than for any given point. (B) Cross-correlation curves c(r) for the examples shown in A. c(r, τ=0) assembles the value of the τ=0 points. These curves do not correct for bleed-through and have large error bounds. c(r, <τ>) averages points tabulated for τ<2s, while c(r, τ→0) extrapolates the slow Gaussian fit to τ=0 (where dashed line intersects y axis in A). These curves represent 2 methods of improving statistics and correcting for bleed-through. These curves are similar, and have lower amplitudes and reduced errors compared to c(r, τ=0).

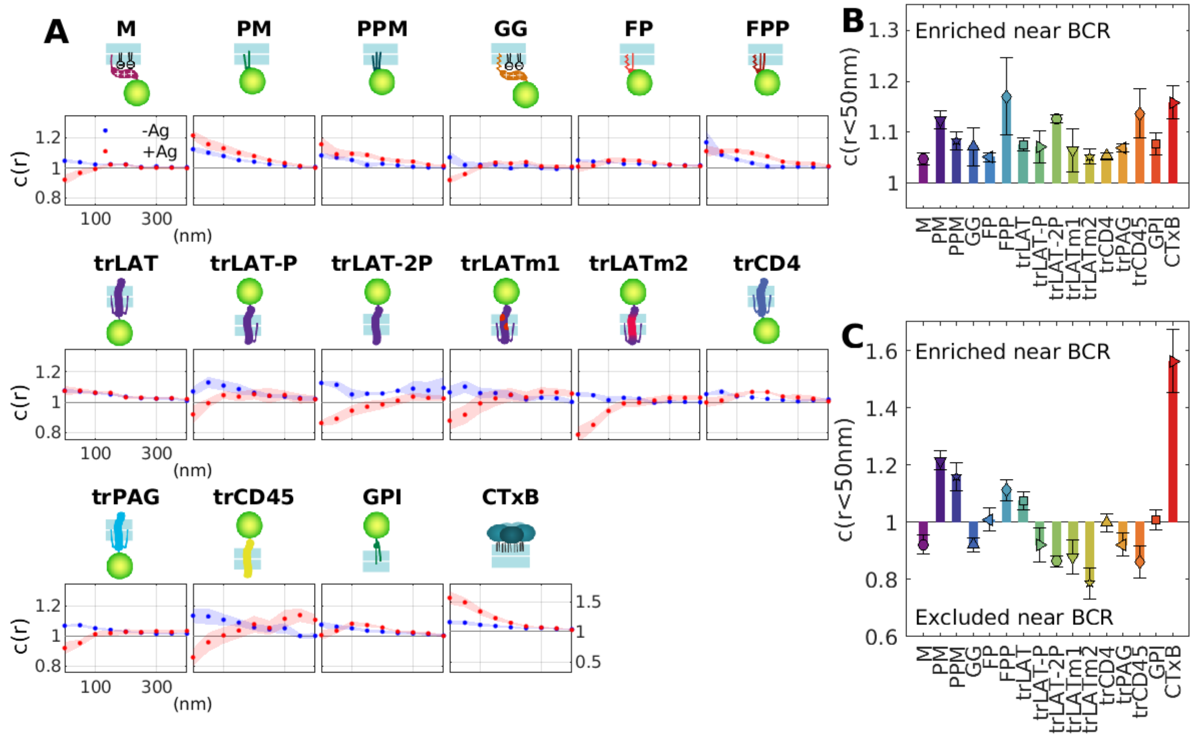

**Supplementary Figure S2: Steady-state cross-correlation functions in cells before and after BCR crosslinking.** (A) Average cross-correlation functions, c(r), between BCR and anchors in cells prior to BCR crosslinking (-Ag) or between 2-10min after streptavidin addition (+Ag). (B,C) Summary showing the cross-correlation values for separation distances less than 50nm from for cells imaged prior to BCR crosslinking (B) or between 2-10min after BCR crosslinking (C). All anchors show weak co-localization with unclustered BCR (c(r<50nm)>1). Amplitudes are more varied across anchors after BCR crosslinking.

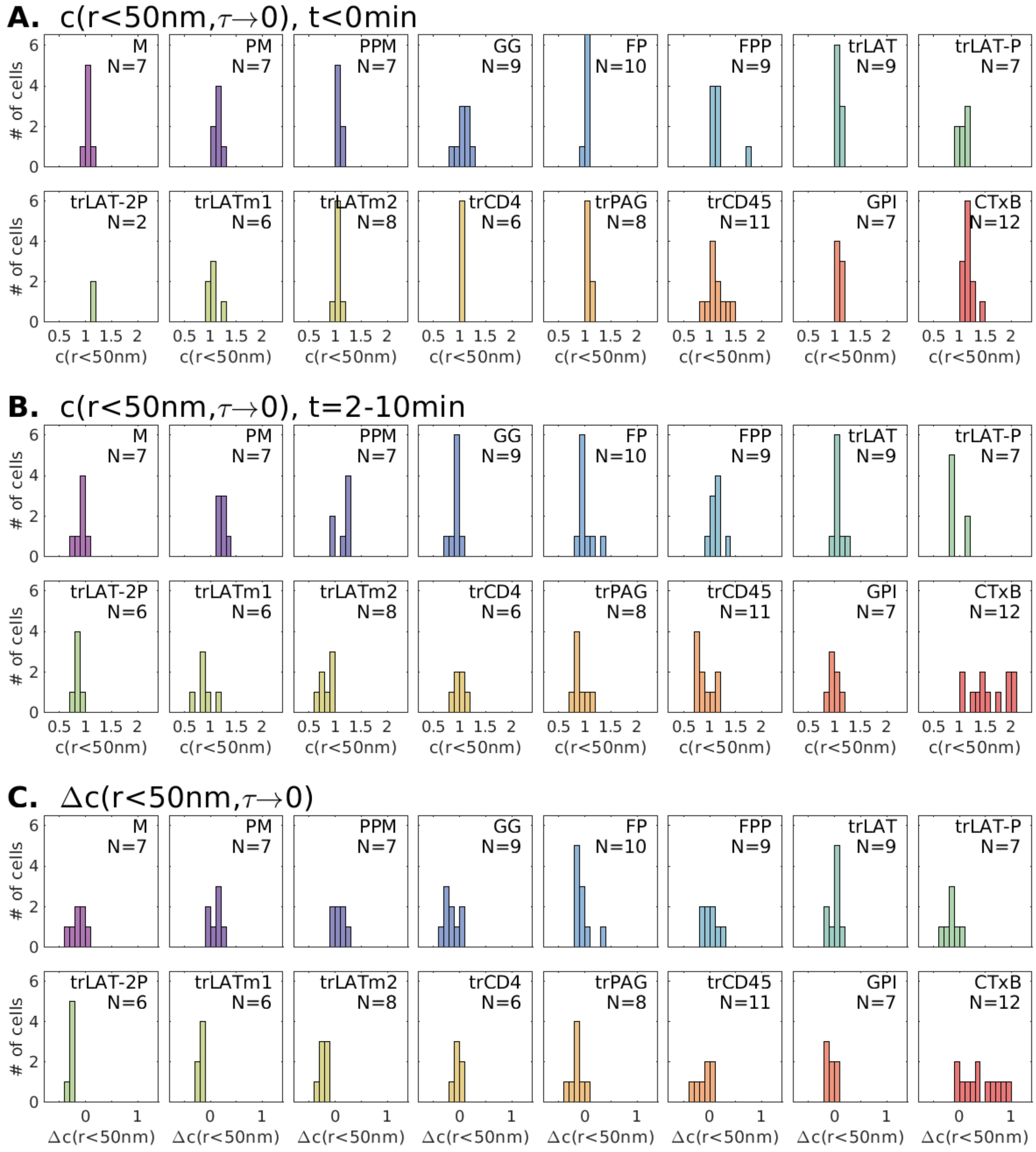


**Supplementary Figure S3: Distribution of corrected cross-correlation amplitudes across cells for all anchors included in this study.** (A,B) Cell-to-cell variation in cross-correlation amplitudes c(r<50nm) tabulated from images acquired prior to BCR clustering (A, t<0min) and from images acquired t=2-10min after BCR clustering (B). (C) Cell-to-cell variation in the change in correlations after crosslinking (Δc(r)) obtained by subtracting values in A from those shown in B. N represents the total number of cells examined for each anchor. For trLAT-2P, not all cells were imaged prior to cross-linking. In this case Δc(r) is obtained by subtracting the average value for the cells expressing this anchor that were imaged prior to BCR clustering.

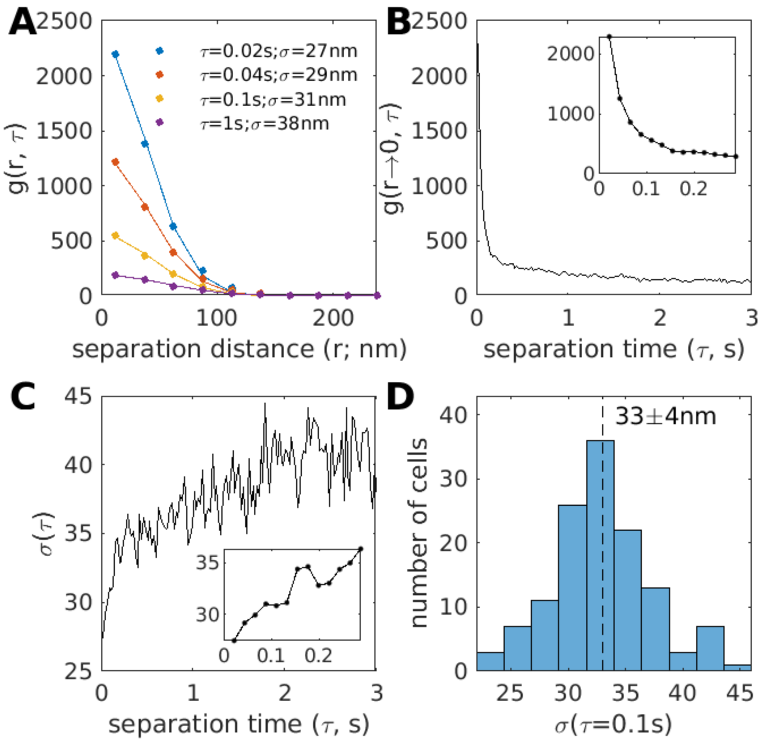


**Supplementary Figure S4: Estimating the size of BCR clusters.** (A) BCR auto-correlation functions (g(r, τ)) for one cell expressing the M anchor between 2-10min after BCR crosslinking. Curves are fit at fixed τ to $g(r, \tau) = 1 + A \times \exp\left\{-\frac{r^2}{4\sigma^2}\right\}$ to extract the amplitude (A) and range (σ) of correlations. (B) Plots of A vs. τ for the same cell. We attribute the large amplitude at short τ (<0.1s) to multiple sequential observations of the same fluorophore, while correlations at larger τ (>0.1s) largely arise from different fluorophores in the same BCR cluster. (C) Plots showing the σ vs τ for the same cell. At short τ (<0.1s), the range of correlations reports on the localization precision of the measurement. At larger τ (>0.1s), σ reflects the size of BCR clusters convoluted with the motion of clusters, which tends to further broaden g(r) at long τ. Based on this we estimate average BCR cluster radius in this cell to be near 34nm, which is roughly σ(0.1s). (D) a histogram showing σ(0.1s) extracted from BCR autocorrelations over all cells is centered at 33nm.

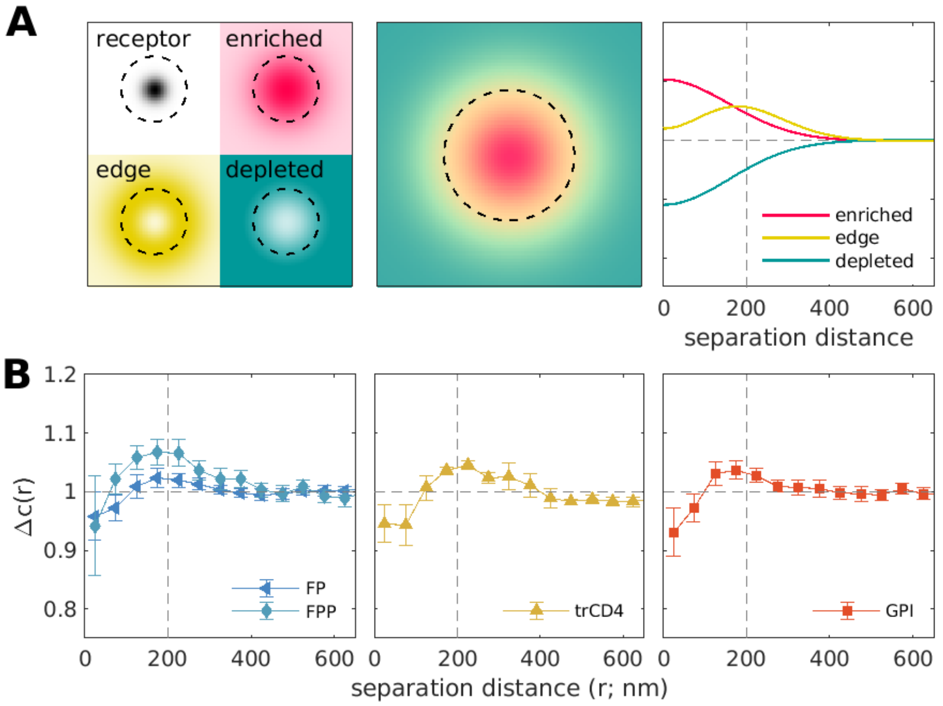


**Supplementary Figure S5: Some anchors enrich at the periphery of BCR clusters.** (A) Schematic showing densities of receptors and hypothetical anchors that are enriched, localized to the periphery of receptor clusters, or depleted from clusters (left) and the resulting shapes of receptor-anchor cross-correlation functions (right). The receptor cluster radius is set based on the BCR cluster size estimated from experiments (Sup Fig S4). Notably the periphery-localized hypothetical anchor gives rise to a cross-correlation function with a local maxima away from r=0 separation distance. Dashed lines represent positions 200nm from the center of the receptor cluster (left) or a separation distance of r=200nm (right). (B) BCR-anchor cross-correlation functions for select inner leaflet peripheral (left), transmembrane (middle), and extracellular (right) anchors that exhibit local maxima in corrected cross-correlation functions for r>0.

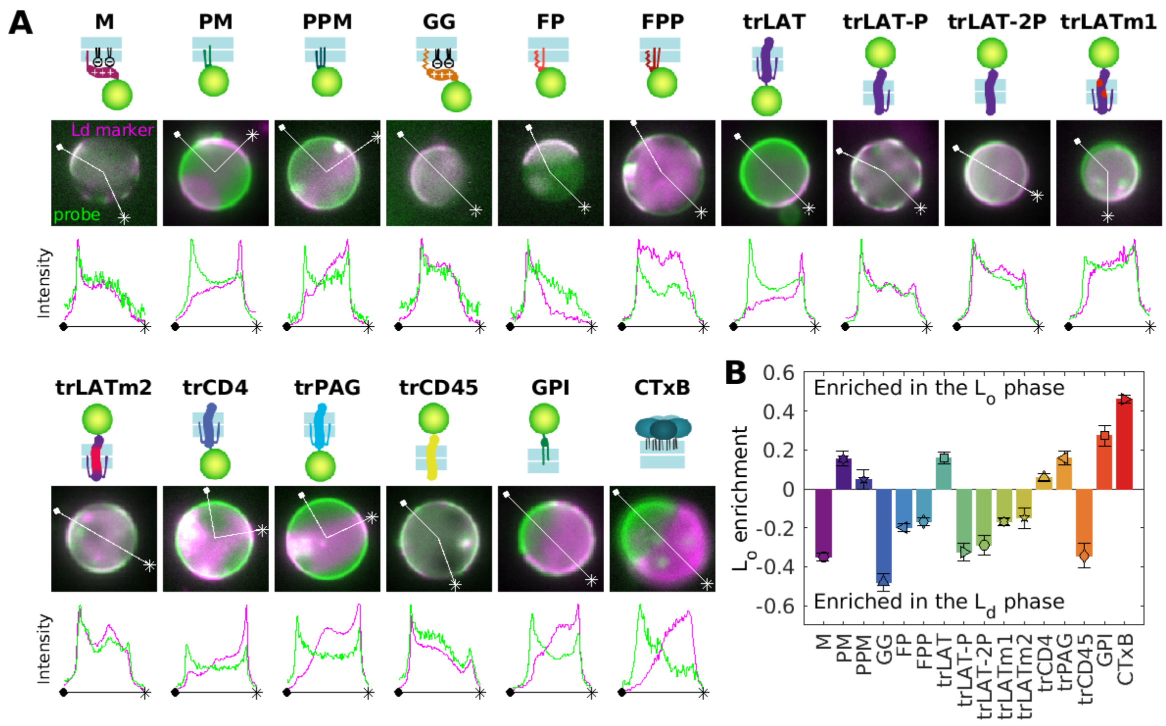


**Supplementary Figure S6: Representative images of GPMVs containing all anchors.** (A) Schematic representation of anchors alongside images of representative GPMVs and intensity traces, as described in Fig 3A. (B) Average values over multiple vesicles and experiments, reproduced from Fig 3B.

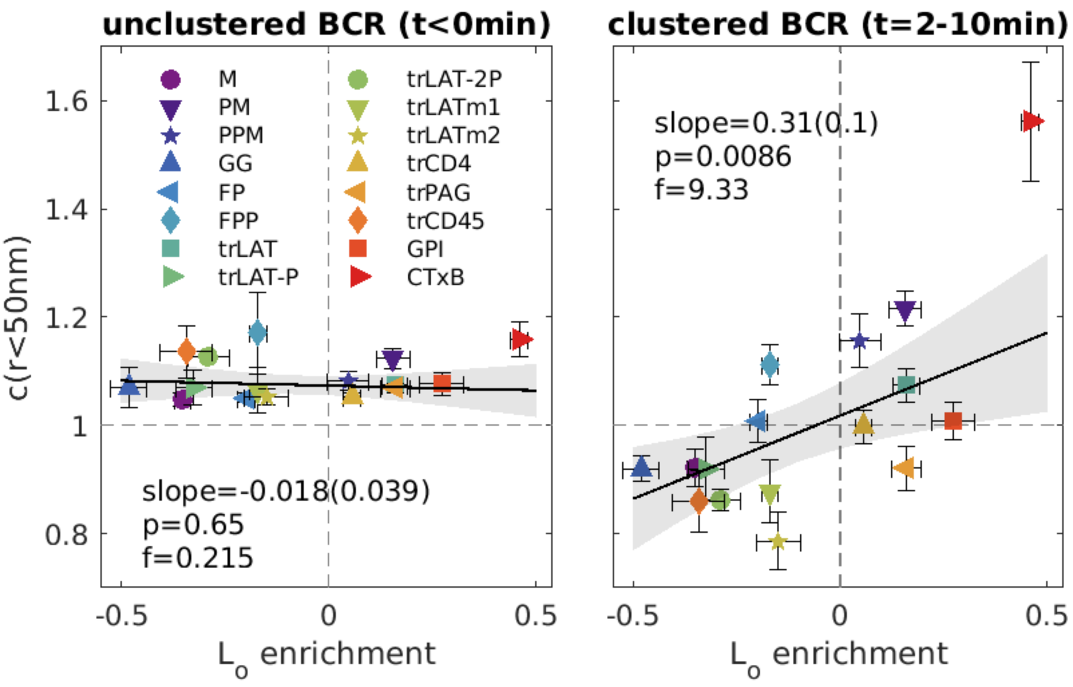


**Supplementary Figure S7: Correlations between GPMV and BCR partitioning of membrane anchors before (left) and after (right) BCR clustering.** Data points are c(r<50nm) values replotted from Supplementary Figure S2 B (left) and C (right) vs Lo enrichment from GPMV measurements (Fig 3B). Trends are fit to a linear model and the significance is assessed with a p-value vs. the null hypothesis of no correlation (lower values indicate greater significance), and an f-statistic that describes how well the variance in the data is described by the linear fit (higher values indicate greater significance). Shaded regions in A-B indicate the 95% confidence interval of the linear fit.

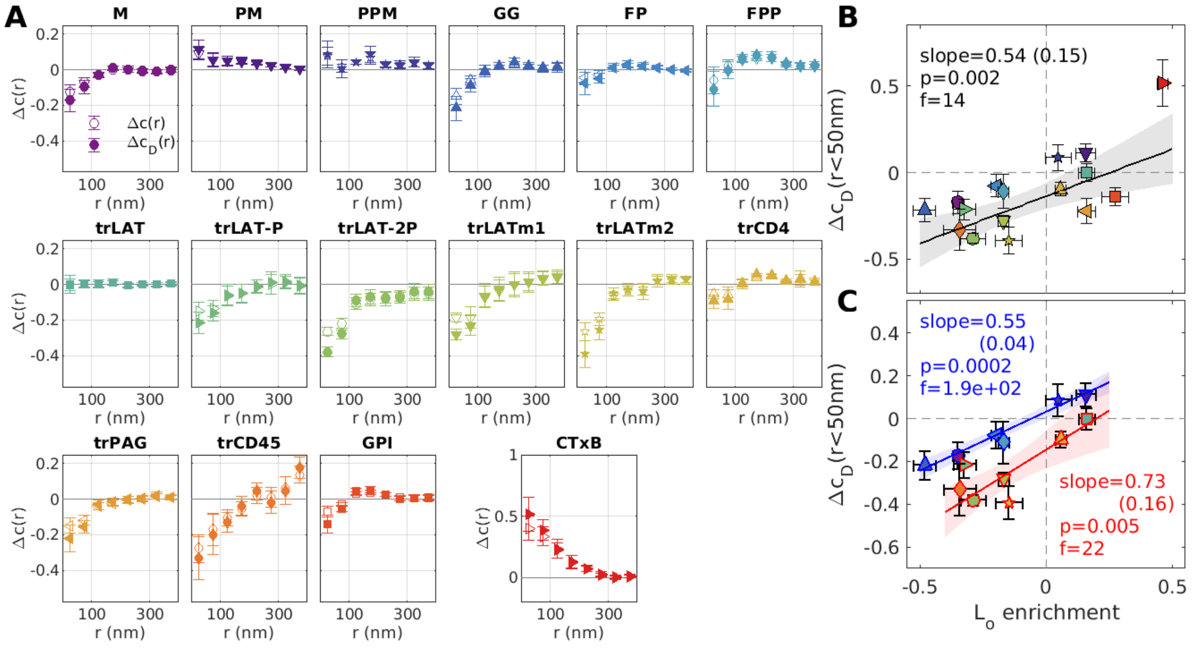


**Supplementary Figure S8: Density enrichment of probes with respect to BCR cluster centers as estimated through deconvolution.** (A) Experimentally determined ($\Delta c(r)$; open symbols) and an estimate of anchor density with respect to BCR cluster centers ($\Delta c_D(r)$; closed symbols) obtained by deconvolving $\Delta c(r)$ with a Gaussian function with a standard deviation of 50nm as described in Methods. (B) $\Delta c_D(r<50nm)$ plotted vs. Lo enrichment in GPMVs, taken from Fig 3B. All points are fit to a linear model with the slope and significance parameters shown. (C) Points representing inner leaflet peripheral anchors (blue edges) and transmembrane anchors (red edge) redrawn from part B but fit to separate linear models (blue and red line respectively) with slopes and significance parameters indicated in colored text.

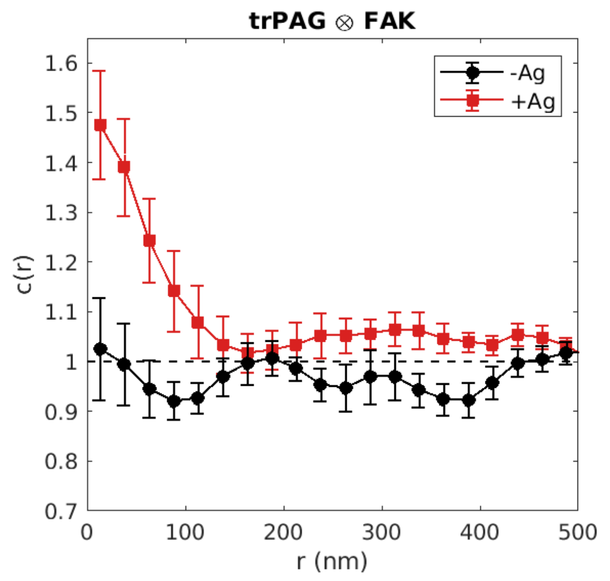


**Supplementary Figure S9: Stimulation dependent reorganization of trPAG with respect to cellular structures labeled by FAK in chemically fixed cells.** CH27 B cells expressing trPAG were chemically fixed either without BCR crosslinking (-Ag) or after 6 min after BCR cross-linking (+Ag). FAK-GFP was expressed in cells and labeled post-fixation with GFP-specific nanobodies as described in Methods.

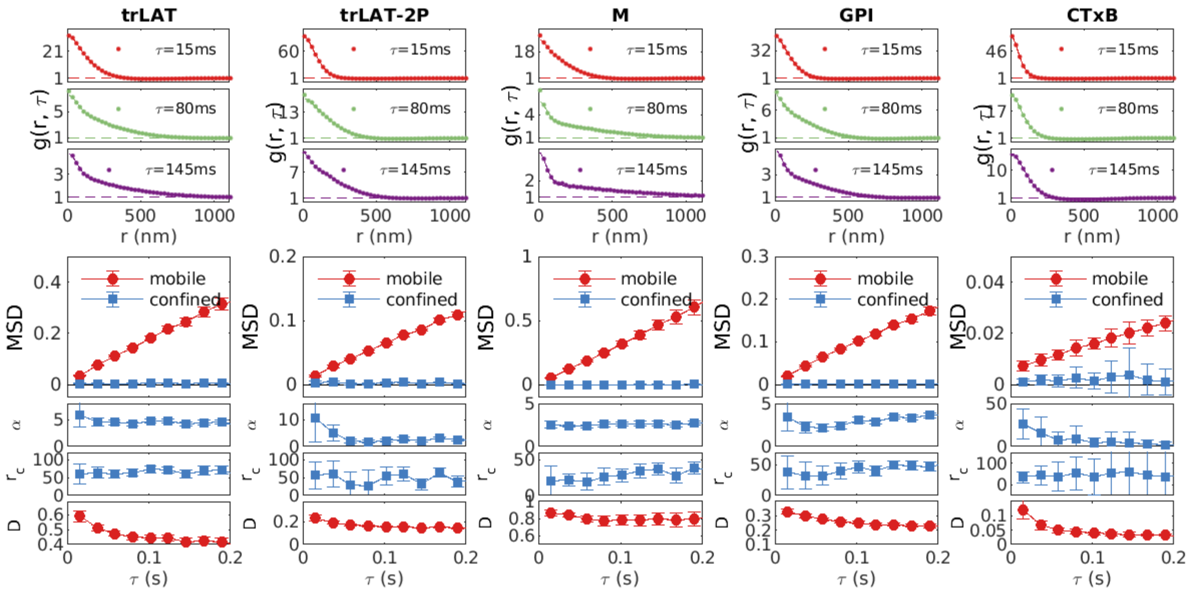


**Supplementary Figure S10: Spatiotemporal auto-correlation functions (g(r,τ)) and extracted fit parameters for several cells and anchors.** For each cell, tabulated points are fit to a superposition of 2 Gaussian functions to extract mean squared displacements (MSDs) and the percentage of molecules in the more confined state (α). The MSD for the confined population can be converted to a confinement radius: $r_c = \sqrt{MSD}$, with units of nm. The MSD for the mobile population is converted to a diffusion coefficient D, with units of $\mu m^2/s$.

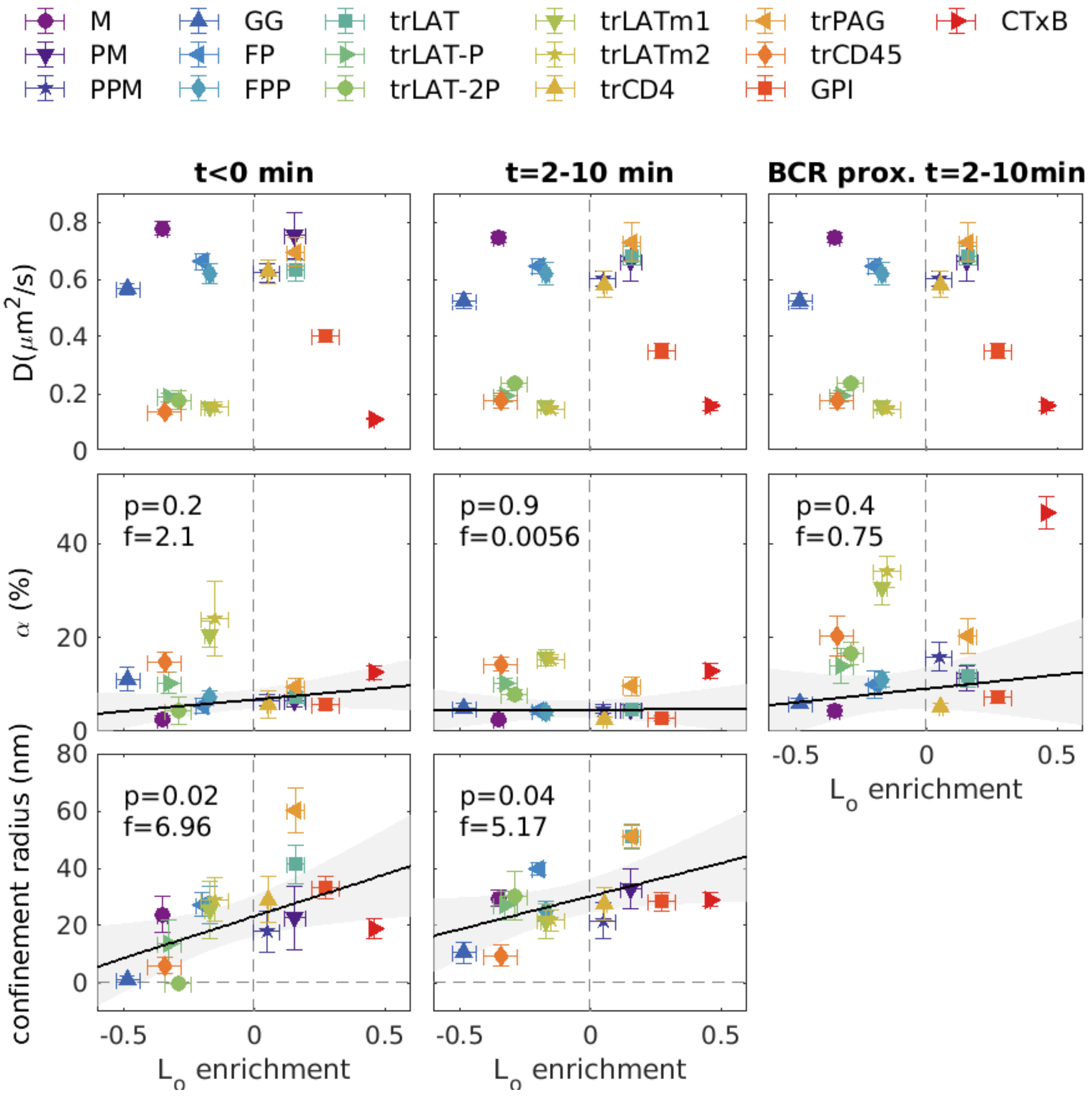


**Supplementary Figure S11: Summary of fit parameters describing anchor mobility at τ=15ms.** Diffusion coefficients (D) for the mobile component, the fraction of molecules in the confined state (α) and the confinement radius averaged over cells expressing the specified anchor for all conditions investigated. Fit parameters for each anchor are plotted vs. Lo enrichment in GPMVs, taken from Fig 3B. The condition "t<0 min" indicates that anchor auto-correlation functions are constructed from images acquired prior to BCR crosslinking and "t=2-10min" indicates that anchor auto-correlation functions are constructed from images acquired between 2 and 10 min after BCR crosslinking. For the condition "BCR proximal", anchor localizations found within 100nm of a BCR localization are cross-correlated with all anchor localizations. The fitting of BCR proximal curves was accomplished by fixing the confinement radius to the value obtained for all trajectories (t=2-10min) to improve the robustness of fitting to curves with lower signal to noise. For confinement radius and α, all points are fit to a linear model with the significance parameters shown.

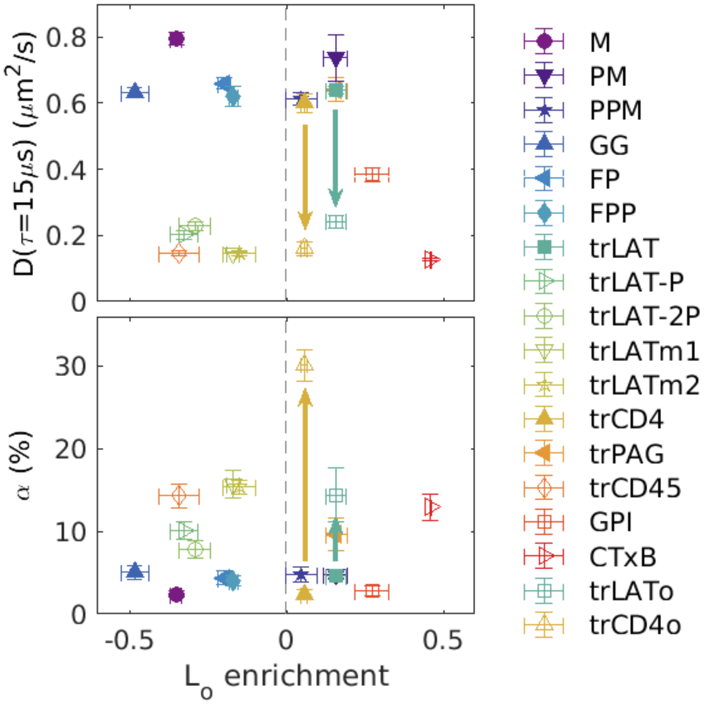


**Supplementary Figure S12: moving mEos3.2 probe to the extracellular terminus slows diffusion (D) and increases the population of confined diffusers (α) for both trLAT and trCD4.** Diffusion coefficients (D) for the mobile component and the fraction of molecules in the confined state (α) are plotted vs. Lo enrichment in GPMVs as in Sup Fig S11. Constructs with extracellular mEos3.2 are shown as open symbols and arrows in the diffusion plot highlight the change for trLAT and trCD4 when the probe is moved from the cytoplasmic side of the transmembrane anchor (trLAT and trCD4) to the extracellular space (trLATo and trCD4o).

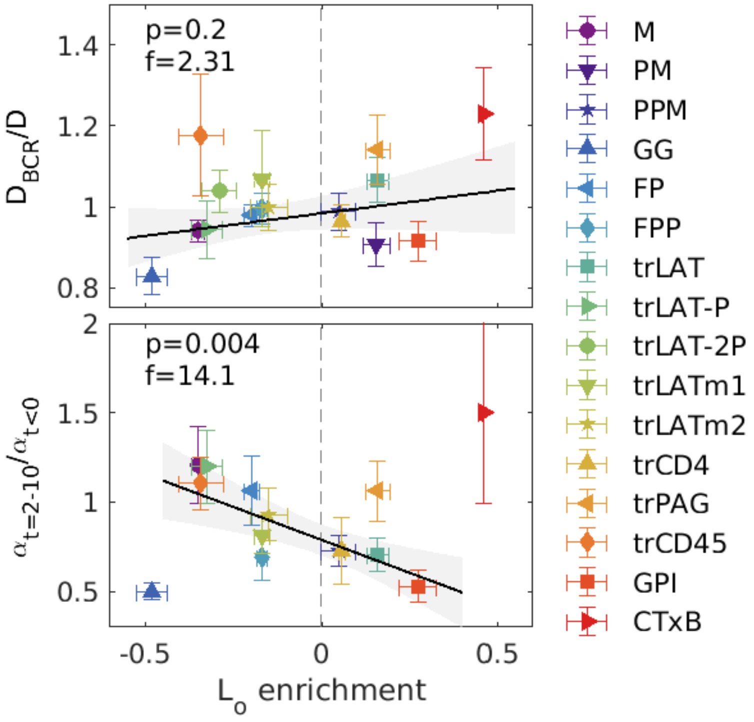


**Supplementary Figure S13: Additional ratios of fit parameters.** (Top) Ratio of the diffusion coefficient of the mobile component for molecules localized near BCR clusters ($D_{BCR}$) to the mobile diffusion coefficient across the entire membrane (D). All points are fit to a linear model with the significance parameters shown and no meaningful trend is observed with Lo enrichment in GPMVs for this ratio. (Bottom) Ratio of the fraction of confined molecules found 2-10 min after BCR crosslinking $\alpha_{t=2\text{-}10}$ to the fraction of confined molecules found prior to BCR crosslinking ($\alpha_{t<0}$). All points other than GG and CTxB are fit to a linear model with the significance parameters shown. A significant trend showing that anchors that enrich in the Lo phase in GPMVs also become less confined after BCR crosslinking. We note that GG is secreted by cells and some immobile molecules visualized early in the experiment (t<0min) are likely adhered directly to the glass coverslip, which could contribute to this anchor representing an outlier here.

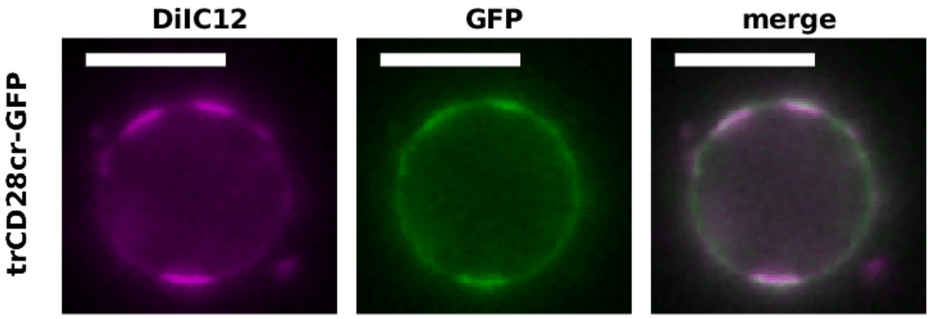


**Supplementary Figure S14: trCD28cr partitions with the Ld phase in GPMVs.** Representative GPMV derived from RBL-2H3 cells expressing trCD28cr-GFP imaged alongside DiIC12, which marks the Ld phase. Scale-bar is 10µm.

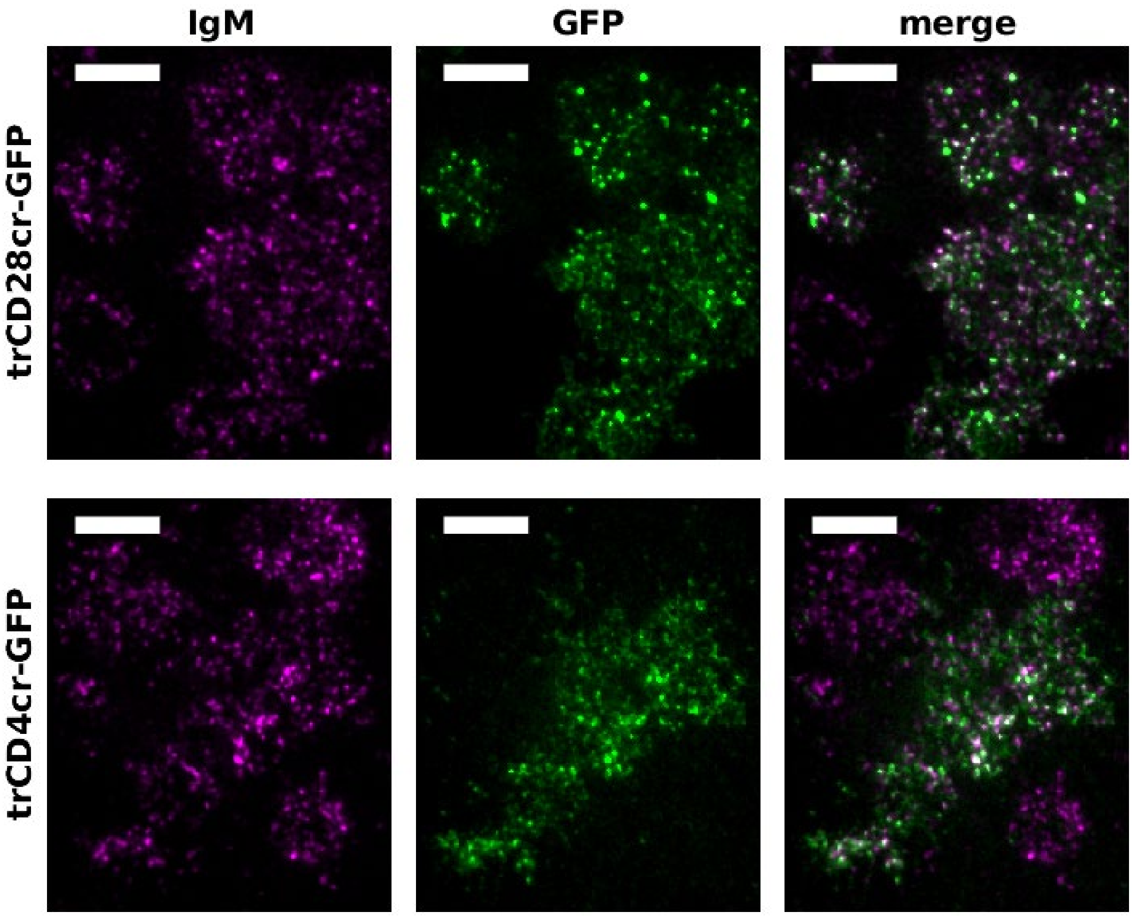


**Supplementary Figure S15: minimal co-receptors co-localize with BCR puncta.** CH27 B cells transduced with either trCD28cr-GFP or trCD4cr-GFP are labeled with biotinylated anti Strep-tagII primary antibodies then co-ligated with biotin-SiR-fAb αIgMμ labeled BCR using streptavidin. Both BCR and minimal co-receptors form puncta that co-loclaize in transduced cells. Individual puncta exhibit a distribution of relative co-receptor and BCR intensities.

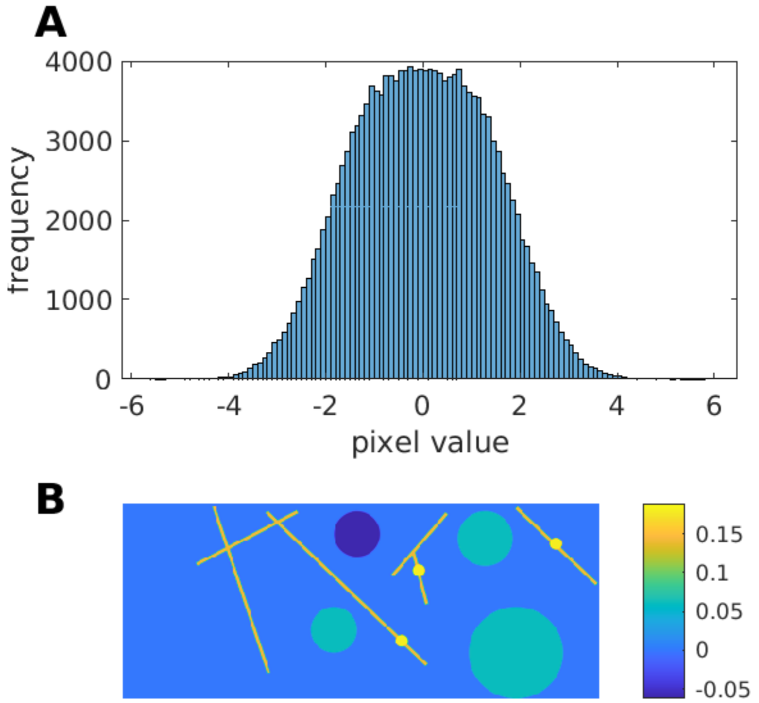


**Supplementary Figure S16: Parameters used to generate the membrane in Figure 7.** (A) The distribution of pixel values used. Half of the values are normally distributed around -1 and half of the values are normally distributed around +1 to create the broad distribution shown. (B) Image representation of the applied Field matrix. Most values are set to 0.


1 Levental I, Lingwood D, Grzybek M, Coskun U, Simons K. Palmitoylation regulates raft affinity for the majority of integral raft proteins. *Proc Natl Acad Sci USA* 2010;**107**:22050–4. https://doi.org/10.1073/pnas.1016184107.

2 Stone MB, Shelby SA, Núñez MF, Wisser K, Veatch SL. Protein sorting by lipid phase-like domains supports emergent signaling function in B lymphocyte plasma membranes. *ELife Sciences* 2017;**6**:e19891. https://doi.org/10.7554/eLife.19891.

3 Núñez MF, Wisser K, Veatch SL. Synergistic factors control kinase–phosphatase organization in B-cells engaged with supported bilayers. *Mol Biol Cell* 2020;**31**:667–82. https://doi.org/10.1091/mbc.E19-09-0507.

4 Lorent JH, Diaz-Rohrer B, Lin X, Spring K, Gorfe AA, Levental KR, *et al.* Structural determinants and functional consequences of protein affinity for membrane rafts. *Nat Commun* 2017;**8**:1219. https://doi.org/10.1038/s41467-017-01328-3.

5 Diaz-Rohrer BB, Levental KR, Simons K, Levental I. Membrane raft association is a determinant of plasma membrane localization. *Proc Natl Acad Sci USA* 2014;**111**:8500–5. https://doi.org/10.1073/pnas.1404582111.

6 Rodgers W. Making membranes green: construction and characterization of GFP-fusion proteins targeted to discrete plasma membrane domains. *Biotechniques* 2002;**32**:1044–6, 1048, 1050–1. https://doi.org/10.2144/02325st05.

7 Pyenta PS, Holowka D, Baird B. Cross-correlation analysis of inner-leaflet-anchored green fluorescent protein co-redistributed with IgE receptors and outer leaflet lipid raft components. *Biophys J* 2001;**80**:2120–32.

8 Pyenta PS, Schwille P, Webb WW, Holowka D, Baird B. Lateral Diffusion of Membrane Lipid-Anchored Probes before and after Aggregation of Cell Surface IgE-Receptors†. *J Phys Chem A* 2003;**107**:8310–8. https://doi.org/10.1021/jp030005t.

9 Ono A, Waheed AA, Freed EO. Depletion of cellular cholesterol inhibits membrane binding and higher-order multimerization of human immunodeficiency virus type 1 Gag. *Virology* 2007;**360**:27–35. https://doi.org/10.1016/j.virol.2006.10.011.

10 Apolloni A, Prior IA, Lindsay M, Parton RG, Hancock JF. H-ras but Not K-ras Traffics to the Plasma Membrane through the Exocytic Pathway. *Mol Cell Biol* 2000;**20**:2475–87.

11 Prior IA, Muncke C, Parton RG, Hancock JF. Direct visualization of Ras proteins in spatially distinct cell surface microdomains. *J Cell Biol* 2003;**160**:165–70. https://doi.org/10.1083/jcb.200209091.
12 Keller P, Toomre D, Díaz E, White J, Simons K. Multicolour imaging of post-Golgi sorting and trafficking in live cells. *Nat Cell Biol* 2001;**3**:140–9. https://doi.org/10.1038/35055042.
